# Intergenerational scaling law determines the precision kinematics of stochastic individual-cell-size homeostasis

**DOI:** 10.1101/2023.01.20.525000

**Authors:** Kunaal Joshi, Charles S. Wright, Rudro R. Biswas, Srividya Iyer-Biswas

## Abstract

Individual bacterial cells grow and divide stochastically. Yet they maintain their characteristic sizes across generations within a tightly controlled range. What rules ensure intergenerational stochastic homeostasis of individual cell sizes? Valuable clues have emerged from high-precision longterm tracking of individual statistically-identical *Caulobacter crescentus* cells as reported in [1, 2]: Intergenerational cell size homeostasis is an inherently stochastic phenomenon, follows Markovian or memory-free dynamics, and cells obey an intergenerational scaling law, which governs the stochastic map characterizing generational sequences of cell sizes. These observed emergent simplicities serve as essential building blocks of the data-informed principled theoretical framework we develop here. Our exact analytic fitting-parameter-free results for the predicted intergenerational stochastic map governing the precision kinematics of cell size homeostasis are remarkably well borne out by experimental data, including extant published data on other microorganisms, *Escherichia coli* and *Bacillus subtilis*. Furthermore, our framework naturally yields the general exact and analytic condition that is necessary and sufficient to ensure that stochastic homeostasis can be achieved and maintained. Significantly, this condition is more stringent than the known heuristic result from quasi-deterministic frameworks. In turn the fully stochastic treatment we present here extends and updates extant frameworks, and highlights the inherently stochastic behaviors of individual cells in homeostasis.

## I . INTRODUCTION

The processes of cellular growth and division are fundamental to the survival and propagation of life. An outstanding open question is how the characteristic generational size of an individual cell remains “constant” across generations, even as that cell repeatedly grows and divides, given the significant stochasticity in both growth and division processes. Homeostasis is the process of maintaining “constancy” to within a specified tolerance of a macroscopic characteristic or state variable, despite the complex internal processes that prevail in even the simplest of organisms [3, 4]. The question of how cells maintain size homeostasis is, in turn, connected to the broader goal of understanding the quantitative principles and mechanisms by which complex processes are controlled and regulated to ensure proper organismal functioning in the face of inescapable stochastic fluctuations in both internal and external environments [5, 6].

Bacterial cells serve as uniquely convenient systems to characterize the dynamics of cell size homeostasis, since the entire organism consists of a single cell. Recently developed approaches for multigenerational singlecell imaging of microorganisms provide the means to observe growth and division tracks of individual bacterial cell sizes, division upon successive division [1, 7–13]. Popular single-cell technologies typically use the concept of the Mother Machine, which takes advantage of designed confinement and one-dimensional arrangement of cells in snug-fitting channels and alleviates the problem of exponential crowding of the imaging region of interest. The “mother” cell confined to the bottom of the narrow channel can be tracked for extended durations since it cannot easily escape and flow away. Thus, these approaches yield long-term intergenerational trajectories of individual cell growth and division; the size of the relevant dataset is set by the number of such mother cells that can be imaged in a microfluidic device at reasonably frequent intervals of time with sufficient spatial resolution to record the size at birth and at division.

Significant effort has been expended in characterizing how the size at division (“final size”) is dependent—on average—on the size at birth (“initial size”) for a given cell in a given generation. Typically, data from different cells and generations are pooled together on a scatter plot of size at division versus size at birth, and the relationship between these quantities used to infer the underlying phenomenology. In this quasi-deterministic picture, homeostasis is characterized by a target “set point” of cell size alongside the sensitivity of final size to initial size in a given generation. This sensitivity is typically categorized into one of three schemes, referred to as sizer (wherein the size at division is independent of the size at birth), timer (wherein the size at division is a multiple of the size at birth), and adder (wherein the size at division is a constant added to the size at birth) [11, 14–2 For cellular homeostasis, this scheme is used to motivate a heuristic argument for a deterministic exponential approach to the target cell size over successive generations [21– The sensitivity value is reproducible in experimental replicates; however, it varies across species, as well as across growth conditions for the same species. That said, the preponderance of current interest rests on the “adder” scheme [22–25]. In contrast, it is well known that the emergence of cell size homeostasis is the result of an inherently stochastic and intergenerational process *for each individual cell*, as we have shown in [1]. Thus, a deterministic exponential relaxation picture motivated by population averages does not faithfully capture observed intergenerational phenomenology. This divergence points to the necessity of developing data-informed accurate stochastic descriptions of intergenerational homeostasis.

In this work we take advantage of recently reported “emergent simplicities” in the stochastic intergenerational size homeostasis of individual cells of *Caulobacter crescentus* [1]. These datasets were obtained using the SChemostat technology, in which the original “mother” cell is retained after each division, and the newborn “daughter” cell leaves the experimental arena; this facilitates high-precision characterization of all cells being imaged [10, 12]. These datasets focus on the same ensemble of statistically identical non-interacting cells over the course of tens of generations, imaged in precisely controlled time invariant growth conditions. In [1], we reported direct evidence of stochastic cell size homeostasis using these data, demonstrating that the intergenerational dynamics are Markovian and that stochastic homeostasis follows an intergenerational scaling law for cell size. In the present work, we build on these experimental observations in *C. crescentus* to develop a data-informed principled theoretical framework, and derive the exact analytic intergenerational stochastic map governing cell size dynamics. We then reanalyze published multigenerational datasets for *Escherichia coli* and *Bacilus subtilis* obtained using the Mother Machine technology [11]. Our exact, analytic results for the predicted intergenerational stochastic map governing the precision kinematics of cell size homeostasis, developed for *C. crescentus* data without fitting parameters [1], are remarkably well borne out by these experimental data on other microorganisms [11]. (See Figs. 2 and 3.) Furthermore, our framework naturally yields the general exact, analytic condition, which is both necessary and sufficient to ensure that stochastic homeostasis can be achieved and maintained (Eq. (17) and Fig. 4). Significantly, this condition is more stringent than the well-known heuristic result for quasi-deterministic frameworks. The fully stochastic treatment we present here extends and updates previous frameworks, and highlights the inherently stochastic behaviors of individual cells in homeostasis.

**FIG. 1.**
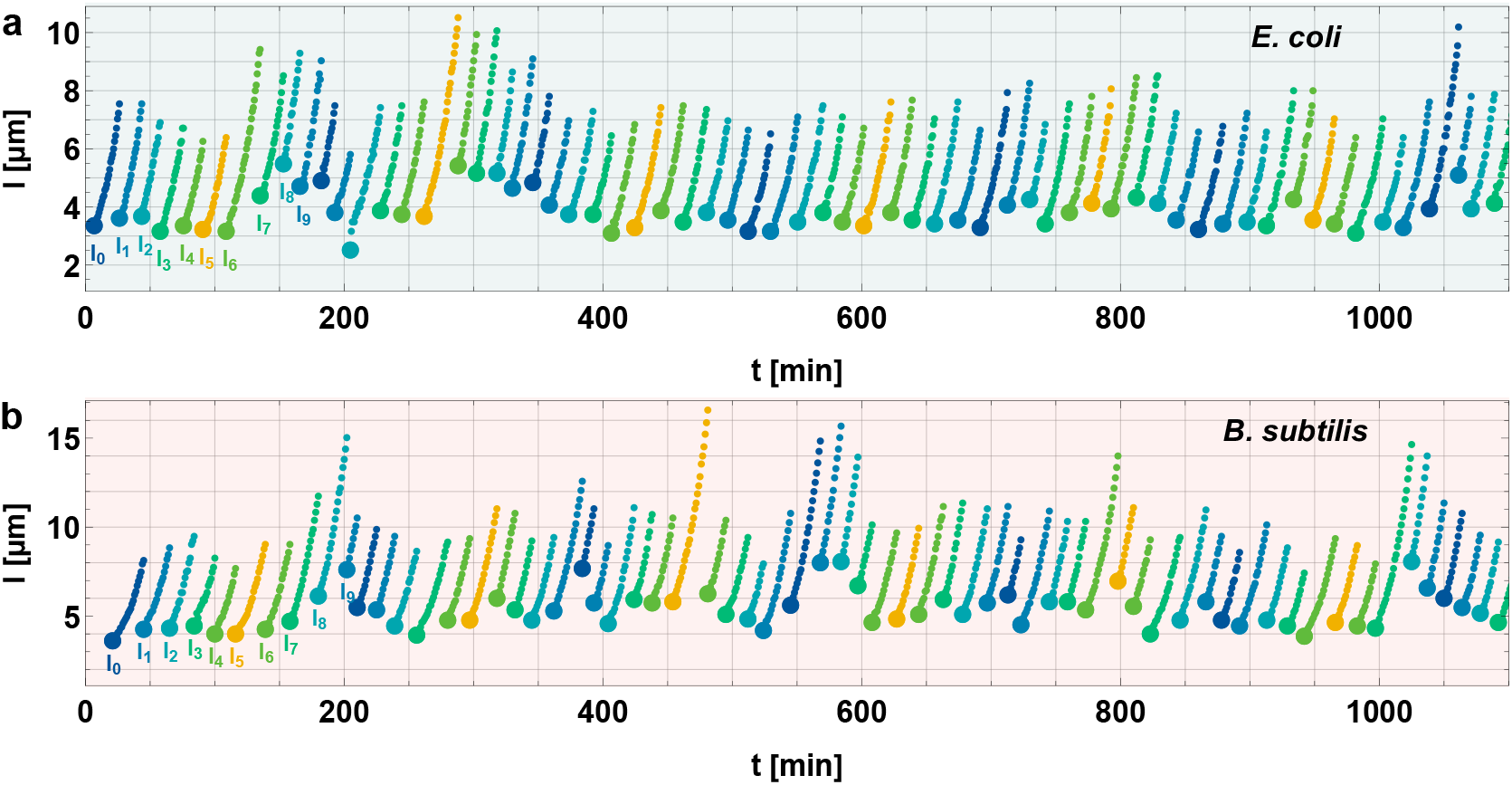
Typical single cell growth trajectories of *E. coli* and *B. subtilis* cells growing in the Mother Machine in steady state. Cell length is plotted as a function of time over multiple division cycles for **(a)** an *E. coli* cell and **(b)** a *B. subtilis* cell grown in TSB media. Cell length was used as a measure of cell size in these datasets. The dimensionful cell lengths at birth, or “initial sizes”, are labeled *l*_*n*_, where *n* is the generation number since the starting, or zeroth, generation. There is considerable variation in initial sizes across generations, yet this variation is bound within a finite range, avoiding runaway sizes. Data source: [11]. (To help make connections with subsequent figures, we note that *a*_*n*_ denotes the initial size in *n*^th^ generation rescaled by population mean initial size.)

**FIG. 2.**
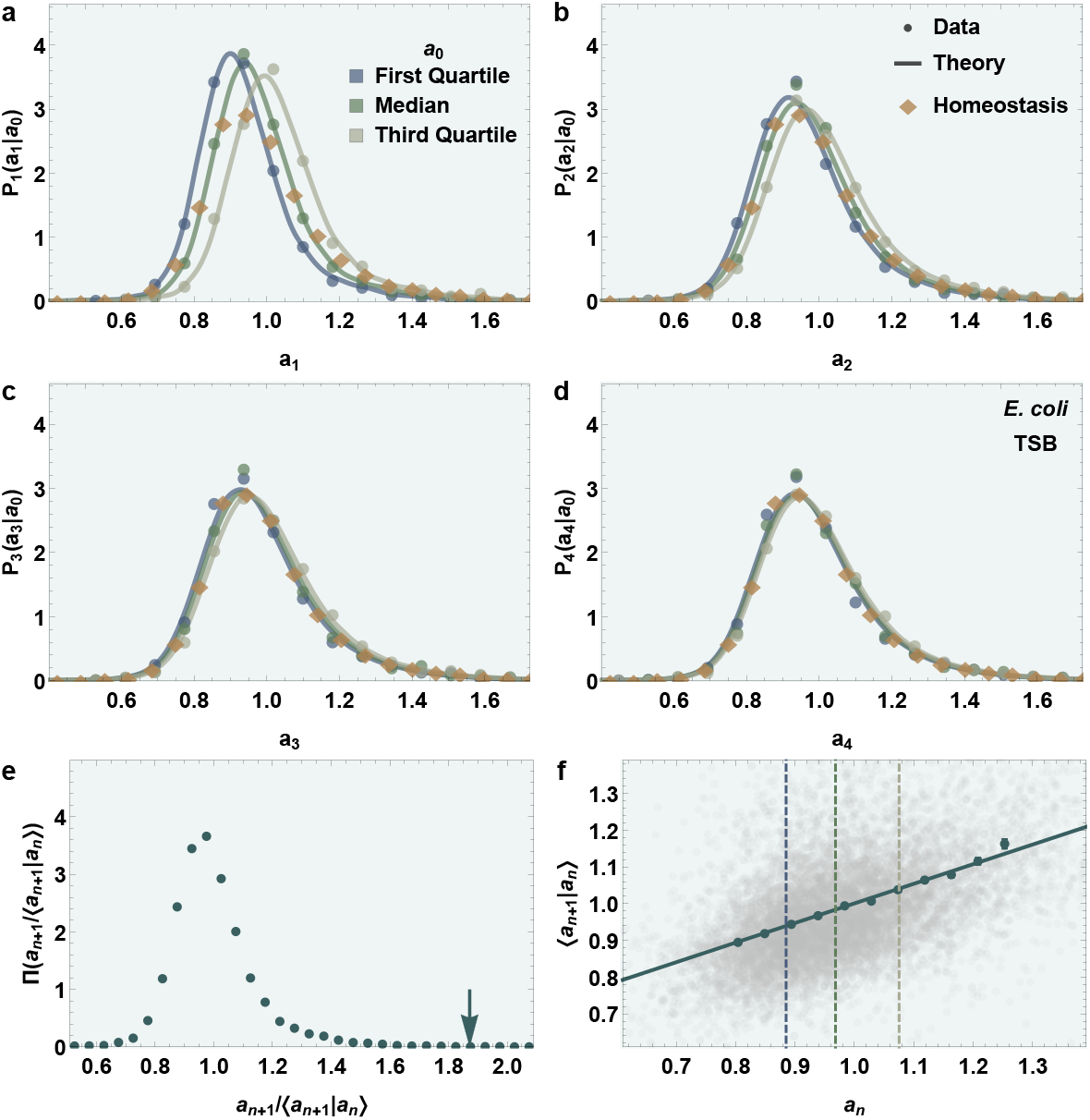
Our theoretical framework accurately predicts, with no fitting parameters, the stochastic intergenerational cell size dynamics leading to homeostasis, for *E. coli* cells. (a–d) Sequence of panels showing the intergenerational evolution of distributions of mean-rescaled initial cell sizes, *a*_*n*_, where *n* is the generation number, starting from three different mean-rescaled initial sizes (consistently chosen to coincide with the first quartile, median, and third quartile for each dataset) in the zeroth generation (*a*_0_, marked by different colors). Panels **(a–d)** correspond to generations 1–4, respectively. The filled circles correspond to experimental data. Curves correspond to fitting-free predictions from our theoretical intergenerational framework. The homeostatic distribution, obtained by pooling together the data for initial sizes, is overlaid in all panels (diamond markers) for ready comparison. Our theoretical framework accurately predicts the stochastic dynamics governing the precise convergence of conditional initial size distributions to the homeostatic distribution over successive generations. **(e–f)** Input functions for predicting initial size distributions over successive generations, and depiction of the necessary and sufficient condition for homeostasis. The experimentally obtained mean-rescaled conditional distribution of the next generation’s initial size, conditioned on the current generation’s initial size, is plotted in (e). The value of 1*/α*, marked by the arrow, agrees with our theoretically derived necessary and sufficient condition for homeostasis: the upper limit of the support of the mean-rescaled distribution should be less than or equal to the corresponding 1*/α*. Here, *α* is the slope of the linear fit of the mean of the next generation’s mean rescaled initial size (*a*_1_), conditioned on the current generation’s mean-rescaled initial size (*a*_0_), plotted in (f). The background scatter plot shows individual data points while the points with error bars show the binned means. The three dashed vertical lines correspond to the three different *a*_0_ of the corresponding color in (a). Data are taken from [11]; these correspond to *E. coli* cells grown in TSB media. Cell length was used as a measure of cell size in these datasets. See Fig. 3 for similar application to *B. subtilis* cells, and the Appendix for application to other growth conditions for both species.

**FIG. 3.**
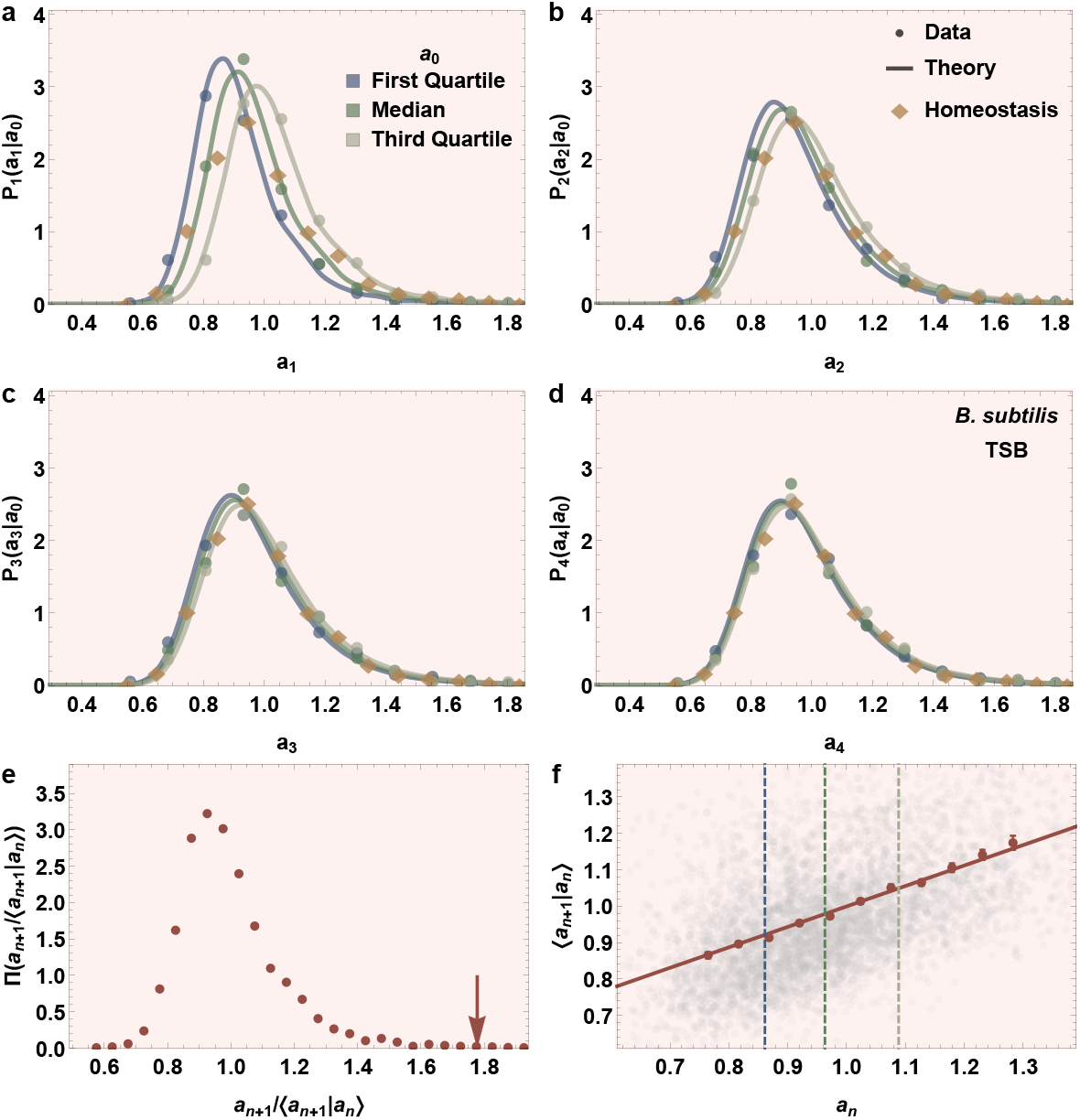
Our theoretical framework accurately predicts, with no fitting parameters, the stochastic intergenerational cell size dynamics leading to homeostasis, for *B. subtilis* cells. (a–d) Sequence of panels showing the intergenerational evolution of distributions of mean rescaled initial cell sizes, *a*_*n*_, where *n* is the generation number, starting from three different mean rescaled initial sizes (consistently chosen to coincide with the first quartile, median, and third quartile for each dataset) in the zeroth generation (*a*_0_, marked by different colors). Panels **(a–d)** correspond to generations 1–4, respectively. The filled circles correspond to experimental data. Curves correspond to fitting-free predictions from our theoretical intergenerational framework. The homeostatic distribution, obtained by pooling together the data for initial sizes, is overlaid in all panels (diamond markers) for ready comparison. Our theoretical framework accurately predicts the stochastic dynamics governing the precise convergence of conditional initial size distributions to the homeostatic distribution over successive generations. **(e–f)** Input functions for predicting initial size distributions over successive generations, and depiction of the necessary and sufficient condition for homeostasis. The experimentally obtained mean rescaled conditional distribution of the next generation’s initial size, conditioned on the current generation’s initial size, is plotted in (e). The value of 1*/α*, marked by the arrow, agrees with our theoretically derived necessary and sufficient condition for homeostasis: the upper limit of the support of the mean-rescaled distribution should be less than or equal to the corresponding 1*/α*. Here, *α* is the slope of the linear fit of the mean of the next generation’s mean rescaled initial size (*a*_1_), conditioned on the current generation’s mean-rescaled initial size (*a*_0_), plotted in (f). The background scatter plot shows individual data points while the points with error bars show the binned means. The three dashed vertical lines correspond to the three different *a*_0_ of the corresponding color in (a). Data are taken from [11]; these correspond to *B. subtilis* cells grown in TSB media. Cell length was used as a measure of cell size in these datasets. See Fig. 2 for similar application to *E. coli* cells, and the Appendix for application to other growth conditions for both species.

**FIG. 4.**
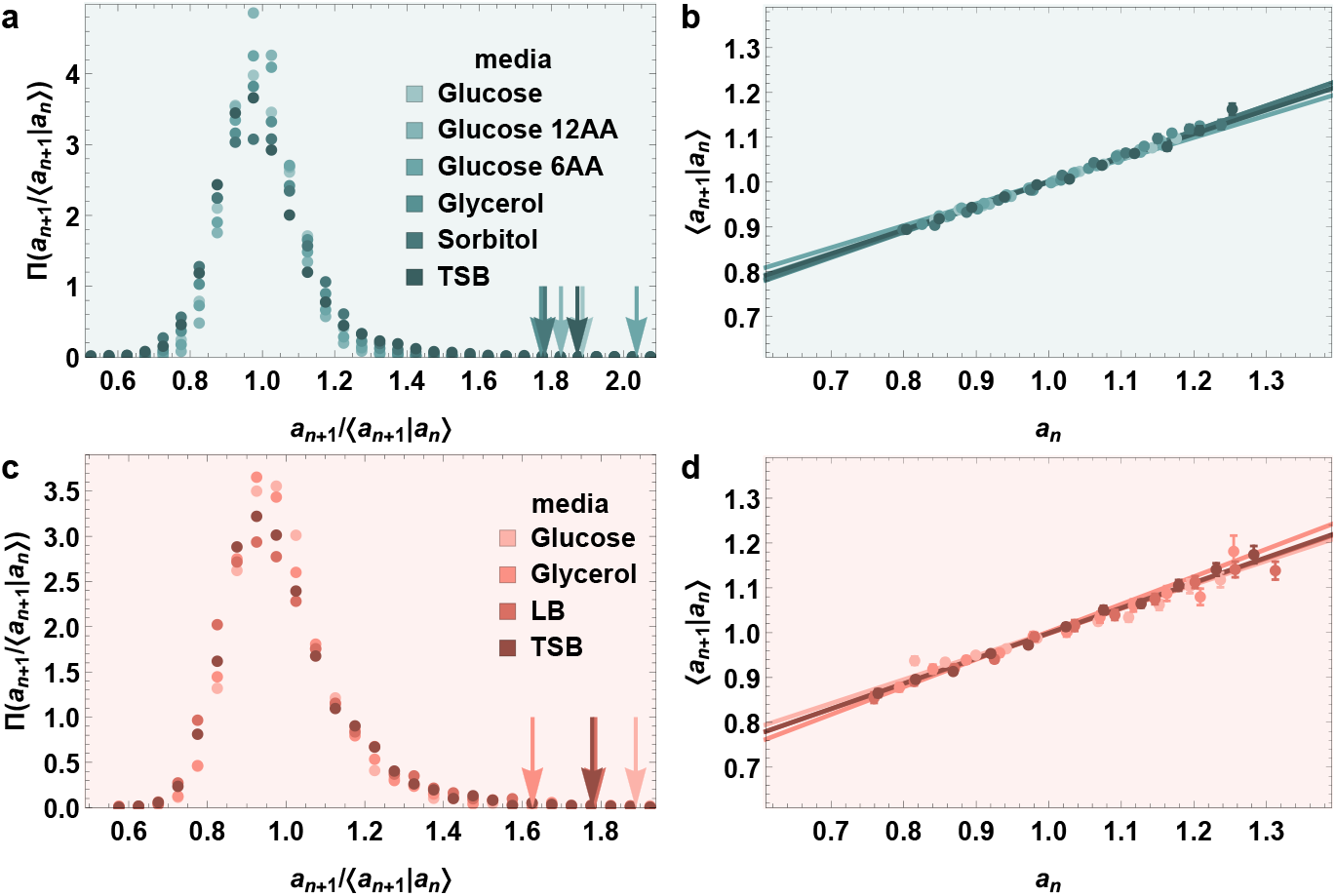
Depiction of the necessary and sufficient condition for homeostasis, and input functions for predicting initial size distributions over successive generations, (a, c) Experimentally obtained mean-rescaled conditional distribution of the next generation’s initial size, conditioned on the current generation’s initial size, for (a) *E. coli* and (c) *B. subtilis* in different growth media (distinguished by color). For each distribution, the value of 1*/α* in the corresponding growth media is marked by an arrow. The arrow placements agree with our theoretically derived necessary and sufficient condition for homeostasis: the upper limit of the support of the mean-rescaled distribution should be less than or equal to the corresponding 1*/α*. Here, *α* is the slope of the corresponding line (with the corresponding color) in (b, d). **(b, d)** Binned means of the next generation’s mean rescaled initial sizes (*a*_1_), conditioned on the current generation’s mean rescaled initial sizes (*a*_0_), are plotted as functions of *a*_0_ along with the corresponding linear fits for the growth conditions with corresponding colors in (a, c). These calibration curves, along with the distributions in (a, c), are used as inputs in our theoretical framework that can then predict the distributions of successive generation’s initial sizes for cells starting from any given initial size (see Fig. 2 and 3). The data used are obtained from [11].

## II. EMERGENT SIMPLICITIES IN THE STOCHASTIC INTERGENERATIONAL HOMEOSTASIS OF INDIVIDUAL CELLS

We now develop a theoretical framework for the generation-to-generation evolution (“kinematics”) of an individual cell’s size (see Fig. I). We use the random variable, *a*_*n*_, to denote the cell size in the *n*^th^ generation. Since cell size grows from division to division, we need to choose a representative size from within the cell cycle: we choose the cell size-at-birth. We also convert to dimensionless units by rescaling all sizes by the populationwide average of the steady state initial sizes. Thus, *a*_*n*_ is the initial size in the *n*^th^ generation, rescaled by the population mean initial size. The motivation for converting cell sizes to dimensionless units (by rescaling by population mean initial sizes) is to unify the results from different experimental setups: In [2], we have shown that for the same species and under the same nutrient conditions, even if the growth and division dynamics in the Mother Machine are different from those in the SChemostat, the kinematics of cell size homeostasis are identical upon rescaling by population mean initial sizes.

We treat *n* = 0 as the “initial” generation. In addition to the usual convention that *P* (*X*|*Y*) denotes the conditional probability distribution of the random variable *X* given a specific realization of the random variable *Y*, we reserve the specific notation *P*_*n*_(*a*|*a*_0_) to denote the conditional probability distribution of *a*_*n*_ at *a*_*n*_ = *a*, given the initial size of the initial generation, *a*_0_. This general setup permits various possibilities for attaining or violating cell size homeostasis, depending on general properties of the multivariate functions *P*_*n*_(*a*|*a*_0_), which in principle can depend on history.

Significant reduction in the complexity of the general problem results from the experimental observations, and corresponding emergent simplicities, for *C. crescentus* cells reported in [1]. The first emergent simplicity is that in each growth condition, the conditional distribution of the next generation’s initial size, conditioned on the current and previous generations’ initial sizes, in fact only depends on the current generation’s initial size. In other words, it is independent of previous generations. Equivalently, the intergenerational dynamics of cell sizes are Markovian under constant growth conditions; thus, the initial size of a given cell in the current generation is the sole determinant of the statistics of future sizes-at-birth of the same cell. It follows that stochastic intergenerational cell size dynamics can be characterized completely by properties of the single-generational stochastic map *P*_1_. In other words, the function *P*_*n*_, for any *n* ≥ 1, can be written down only in terms of *P*_1_:

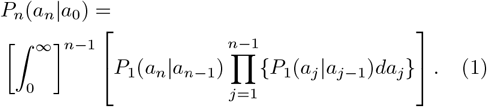

Within this perspective, attainment of cell size homeostasis corresponds to:

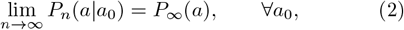

where *P*_∞_ is the asymptotic homeostatic distribution, independent of the initial size. Furthermore, we require that the homeostatic distribution have well-defined moments. Thus, for a cell that starts with initial size *a*_0_, considering all possible futures after a large number of generations, the size probability distribution converges to a well-behaved homeostatic distribution which is independent of the precise value of *a*_0_. In this work, we seek to define under what conditions the kinematics of intergenerational cell sizes specified by the experimentally observed emergent simplicities lead to the homeostatic distribution, *P*_∞_.

In [1] we report a second emergent simplicity for *C. crescentus* cells: the mean-rescaled distribution of cell sizes-at-birth (initial sizes) in a given generation, conditioned on the initial size of the previous generation, is independent of the previous generation’s initial size. We refer to this result as the “intergenerational scaling law”. Mathematically,

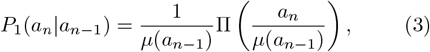

where Π is the universal mean-rescaled distribution that is independent of a cell’s history of size dynamics. Π has mean *m*_1_ = 1, and *µ*(*a*_*n*−1_) is the mean of *a*_*n*_ when conditioned on the previous generation’s initial size, *a*_*n*−1_. We note that the shape of Π may vary depending on growth condition.

Experimentally, *µ*(*a*) is found to be well-described by a linear function over the physiologically relevant range of cell sizes observed (see Fig. 4 B and [1]):

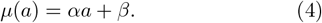

The different values of the slope, *α* (observed in different growth conditions for different organisms), have been scrutinized in great detail, leading to vigorous debates regarding their mechanistic implications for growth control [11, 22–25]. The specific values to which meanings are attached are *α* = 0 (sizer), *α* = 1 (timer), and *α* = *r* (adder, with *r <* 1 being the division ratio). As indicated previously, these approaches view cell size homeostasis as a quasi-deterministic process in which the size in a given generation, if different from the homeostatic setpoint, exponentially relaxes to the setpoint size over successive generations with rate ln(1*/* |*α*|) (see [21]). Since this rate must be positive for homeostasis to be achieved, this heuristic treatment requires that the condition |*α*| *<* 1 be obeyed to permit the possibility of homeostasis. Thus, the value of *α* needs to be less than the slope of the timer (*α* = 1) for homeostasis to be attainable in this quasi-deterministic picture.

In contrast to the quasi-deterministic perspective summarized above, our main focus here is on the intergenerational scaling law, Eq. (3)—we view the value of the slope, *α*, in Eq. (4) as simply a calibration parameter, and do not imbue it with any special mechanistic meaning. We now proceed to develop the precise kinematics governing stochastic intergenerational cell size dynamics. We use the intergenerational scaling law Eq. (3) and the calibration curve Eq. (4) as the essential building blocks of our theory.

## III. RESULTS AND DISCUSSIONS

### A. The intergenerational scaling law universally determines the precision kinematics of stochastic intergenerational homeostasis

We now proceed to develop the following universal framework for intergenerational evolution of an individual cell’s size. Eq. (3) and the Markov property imply that the sizes-at-birth of a given cell follow the stochastic map:

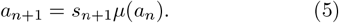

In the preceding equation, the rescaled random variables, {*s*_*n*_}, are uncorrelated. They are drawn independently from the mean-rescaled probability distribution, Π, introduced in Eq. (3), which has mean *m*_1_ = 1. (We denote the *k*^th^ moment of the probability distribution Π by *m*_*k*_.) The random variable *a*_*n*_ denotes the initial cell size in the *n*^th^ generation. Using Eq. (5) recursively, the size-at-birth in the *n*^th^ generation can be related to the size-at-birth in the 0^th^ generation:

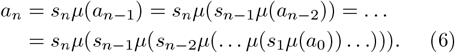

Homeostasis is attained when the probability distribution for *a*_*n*_, and hence all moments of *a*_*n*_, reach finite asymptotic limits that are independent of *a*_0_ as *n* → ∞.

For experimentally relevant scenarios *µ*(*a*) is typically a linear function, *µ*(*a*) = *αa* + *β*, as written down in Eq. (4). Thus Eq. (6) becomes:

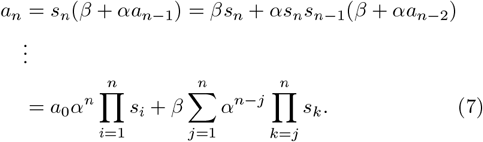

This equation shows the explicit connection between the starting size, *a*_0_; the stochastic variable denoting the size in the *n*^th^ generation, *a*_*n*_; and the sequence of intermediate independent stochastic scaling factors {*s*_1_, *s*_2_, *…, s*_*n*_}.

Equipped with this map, we use our theoretical framework to predict the distributions of initial sizes over successive generations, for cells starting from any given initial size, using only the mean-rescaled distribution Π and the calibration curve *µ*. Our fitting parameter-free predictions (Eqs. (1), (3) and (4)) match excellently when applied to published *E. coli* (Fig. 2) and *B. subtilis* (Fig. 3) data, taken from [11]. Moreover, compelling data-theory matches were also obtained for the *C. crescentus* data as detailed in [1]. Thus we have comprehensively validated the framework we propose here.

### B. The deterministic limit of stochastic intergenerational homeostasis

In what follows, we take *a*_0_ to have a fixed value. All averages ⟨ ⃜⟩ are taken with respect to the random variables {*s*_*m*_}. When *a*_0_ is explicitly taken to be random, i.e., the initial generation size is drawn from an arbitrary distribution, we denote further averaging over initial sizes by double angular brackets, ⟨⟨ ⃜⟩⟩. To evaluate the moments characterizing the probability distribution of *a*_*n*_ in terms of *α, β* and the raw moments, {*m*_*k*_}, of the generation-independent scaling factor distribution Π, we raise Eq. (7) to different powers and take averages. Directly averaging Eq. (7), we find the mean initial size in the *n*^th^ generation:

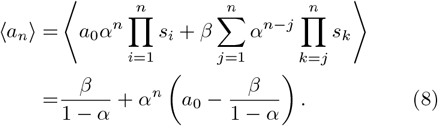

After a large number of generations have elapsed (when *n* → ∞), the second term diverges unless |*α*| *<* 1. *Thus* |*α*| *<* 1 *is a necessary condition for homeostasis*. When |*α*| *<* 1:

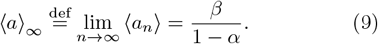

Thus, the asymptotic mean, ⟨*a*⟩_∞_, also becomes independent of *a*_0_ as *n*→ ∞ when |*α*| *<* 1, showing that *the condition* |*α*| *<* 1 *is also sufficient for homeostasis at the level of the mean*.

The deviation of the population mean from its asymptotic (homeostatic setpoint) value decreases exponentially over successive generations, with the exponential rate given by ln(1*/α*). To see this, we use Eq. (9) in Eq. (8) and average over the probability distribution of *a*_0_:

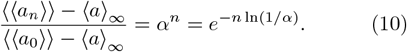

The preceding result from the fully stochastic picture presented here is reminiscent of the deterministic heuristic picture sketched to motivate how homeostasis is attained in the sizer–timer–adder framework (for an example, see Fig. 1 in [21]).

### C. Beyond the mean: fluctuations impose additional requirements for attainment of stochastic homeostasis

To include the effects of fluctuations, we find the variance of size in the *n*^th^ generation by subtracting Eq. (8) from Eq. (7), and squaring and averaging the result:

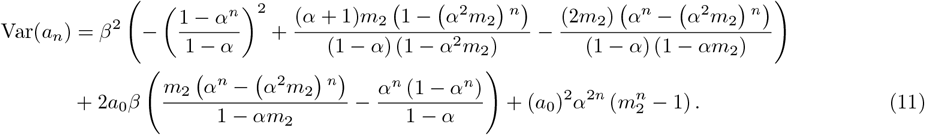

This result implies an additional condition that must be met for homeostasis of the variance, beyond the condition |*α*| *<* 1 required for homeostasis of the mean. That additional condition is |*α*^2^*m*_2_| *<* 1, i.e., |*α*| *<* 1*/ m*_2_, needed for achieving both *a*_0_-independence and a finite limit for Var(*a*_*n*_) at large *n*:

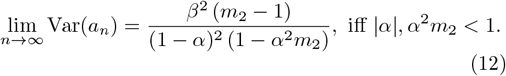

Comparing Eqs. (11) and (12), we note that the variance’s approach to its asymptotic value is through a superposition of exponentials with rates given by ln(*α*), 2 ln(*α*) and ln(*α*^2^*m*_2_). Thus, it is not a simple exponential as in the case of the mean (Eq. (10)).

Since *m*_2_ ≥ (*m*_1_)^2^ = 1, the new condition 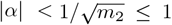 is more restrictive than |*α*| *<* 1. Therefore we have obtained a new condition on cell size homeostasis, which directly results from including the precise intergenerational stochastic behavior of cell growth and division.

**TABLE 1.**
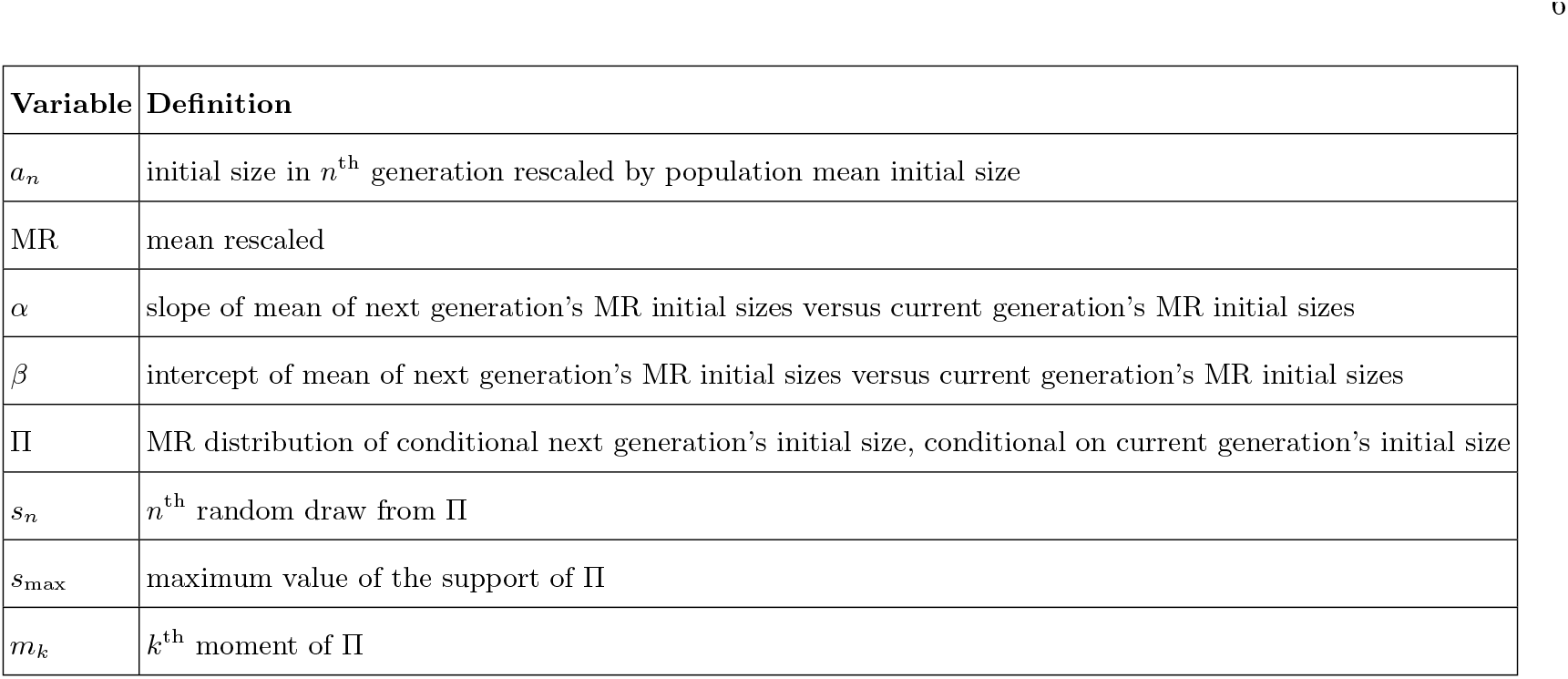
List of variables.

### D. General necessary and sufficient conditions for stochastic intergenerational homeostasis

Our analysis of the asymptotic behavior of all raw moments yields the following general constraints for achieving size homeostasis, when *α* ≥ 0:

For the *k*^th^ moment of the initial cell size to display homeostasis (i.e., it has a finite *a*_0_-independent limit after a large number of generations), all quantities {|*α*^*r*^*m*_*r*_|}, for 1 ≤ *r* ≤ *k*, need to be less than 1. In particular, since *m*_1_ = 1, the first of these conditions is simply |*α*| *<* 1, as can be derived in the deterministic picture. For size homeostasis to hold, however, all moments of the cell size need to display finite *a*_0_-independent asymptotic behavior as the number of generations grows, thus requiring |*α*^*k*^*m*_*k*_| *<* 1 for all *k* ≥ 1.

We derive these constraints, and also an exact expression for the general raw moment of *a*_*n*_, in the Appendix. There we also extend our analysis to *α <* 0, showing an additional constraint that needs to be met for homeostasis:

When *α <* 0, for size homeostasis to hold (i.e., all moments of the cell size display finite *a*_0_-independent asymptotic behaviors as the number of generations grows and *a*_*n*_ never becomes negative) we require |*α*^*k*^*m*_*k*_| *<* 1 for all *k* ≥ 1, and 0 *< a*_0_ *< β/*|*α*|.

Furthermore, when the above conditions are met, the asymptotic values of the raw moments of *a*_*n*_ are:

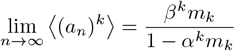

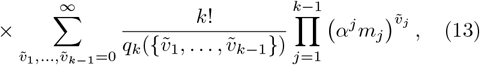

where 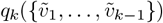 equals one times (*l*+1)! for each set of consecutive zeros of length *l* in the sequence 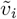’s (see Appendix for further details). As an application, we now demonstrate the equivalence of this formula to Eqs. (9) and (12) (cases *k* = 1 and 2).

Setting *k* = 1 in Eq. (13), the summation is absent and the formula simplifies to Eq. (9) (recall: *m*_1_ = 1). To compare with the *k* = 2 case, first we calculate the second raw moment from Eqs. (9) and (12) :

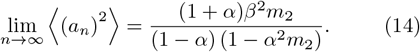

Then, setting *k* = 2 in Eq. (13) and using *m*_1_ = 1:

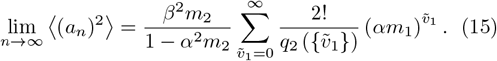

Now, 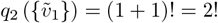 when 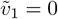 (a ‘string’ of zeros with length 1), and 1 otherwise. Using this,

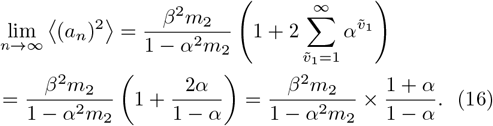

This is the same as Eq. (14), demonstrating their equivalence.

In Fig. 6 we show the generational evolution of the first two moments of the cell size distribution, as the value of *α* is varied through successive homeostasis thresholds, showing successive levels of breakdown in homeostasis.

**FIG. 5.**
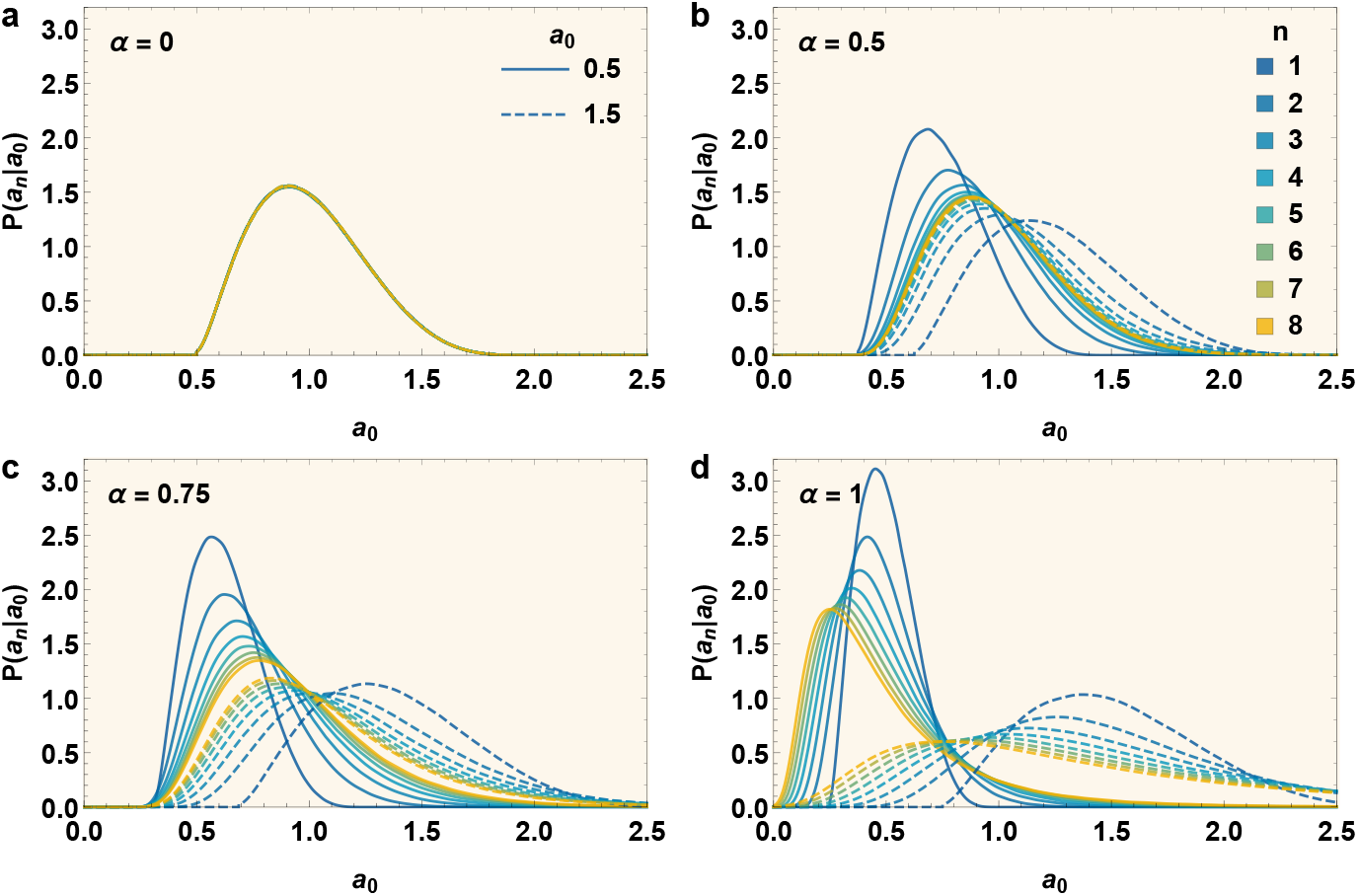
In-silico evolution of conditional initial size distributions over successive generations, for different values of *α*, for cells starting out with a given initial size. These plots are made using the inter-generational kinematic theory presented in Eqs. (1), (3) and (4). Each plot shows the conditional initial size distributions over successive generations (different colors) for cells starting from two different initial sizes, *a*_0_ = 0.5 (solid line) and 1.5 (dashed). 1*/s*_*max*_ = 0.5 for our hypothetical mean-rescaled distribution, given by the shifted Beta distribution Π(*s*) = (2*/*3)^8.5^[Γ(7.5)*/*(Γ(5)Γ(2.5))](*s* −0.5)^2.5^(2 −*s*)^5^ for 0.5 ≤ *s* ≤ 2. The mean of the next generation’s initial size given the current generation’s is ⟨*a*_1_ |*a*_0_⟩ = *µ*(*a*_0_) = *αa*_0_ + (1 −*α*), ensuring that the homeostatic mean setpoint is 1 and allowing easy comparison between different *α*’s. **(a)** *α* = 0. Cells reach the homeostatic distribution within a generation. This corresponds to the sizer model in the deterministic picture. **(b)** *α* = 0.5. Cells reach the homeostatic distribution less rapidly than in (a). Since this value of *α* also satisfies the condition for homeostasis (*α* ≤ 1*/s*_*max*_), all moments are expected to converge to finite values. This corresponds to the adder model in the deterministic picture. **(c)** *α* = 0.75. The condition for homeostasis is no longer satisfied. The mean still converges, and thus the overall location and shape of the distribution appears to converge. However, the higher moments diverge, manifesting in the form of a long tail of the distribution. In the purely deterministic picture, this value of *α* would successfully result in homeostasis, but not in the exact stochastic framework we present here. **(d)** *α* = 1. Here, even the mean does not converge to an *a*_0_-independent value. This corresponds to the the timer model. Homeostasis is visibly absent.

**FIG. 6.**
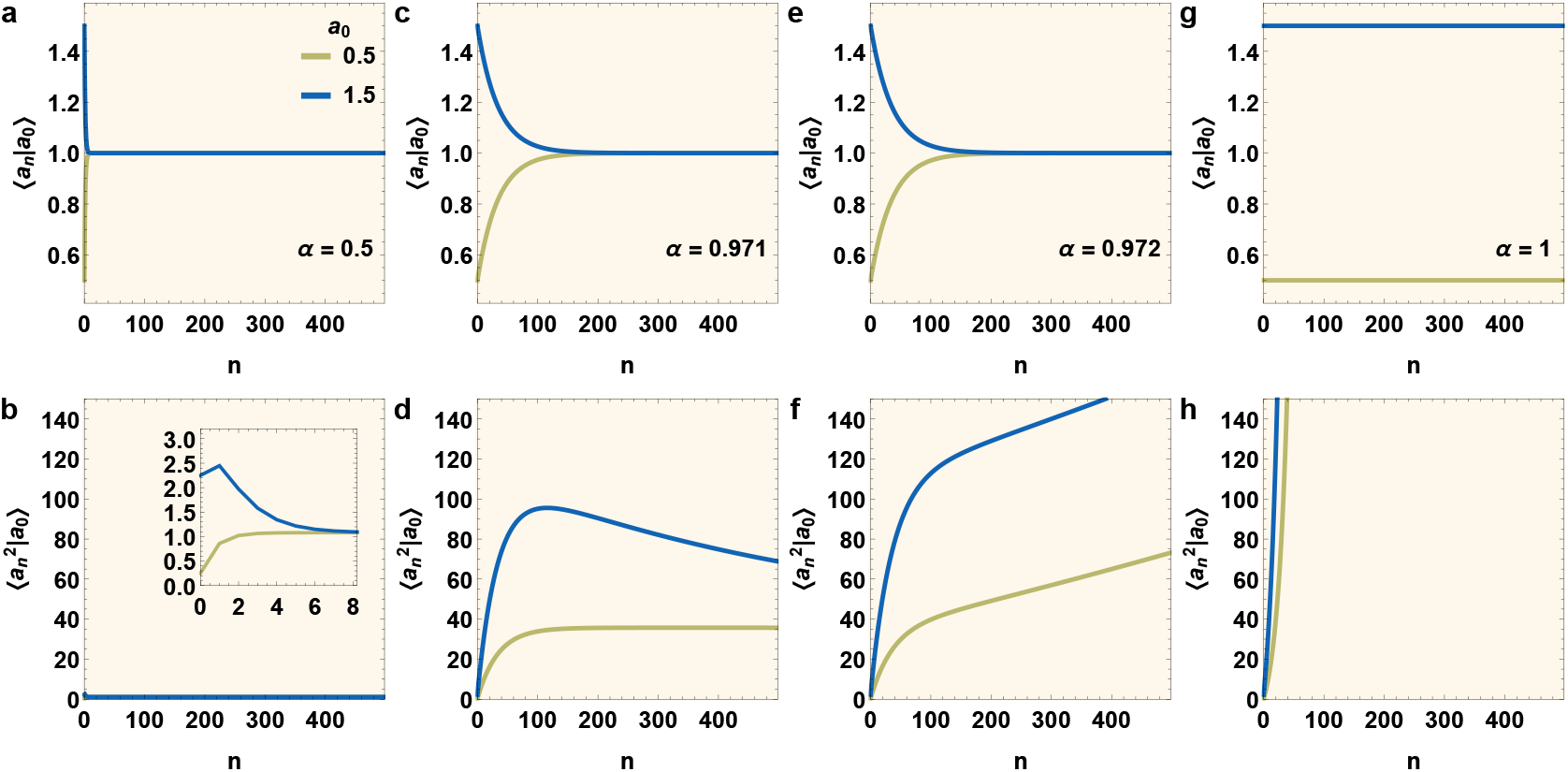
Demonstrating the conditions for convergence of moments. Plots made using the expressions in Eqs. (8) and (11). Each plot shows the evolution of the mean (top panels) or the second moment (bottom panels) of the conditional initial size distributions with generation *n*, for cells starting from two different initial sizes, *a*_0_ = 0.5 (olive) and 1.5 (blue). 1*/s*_*max*_ = 2 and 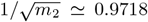 for our hypothetical mean-rescaled distribution, given by the shifted Beta distribution Π(*s*) = (2*/*3)^8.5^[Γ(7.5)*/*(Γ(5)Γ(2.5))](*s* 0.5)^2.5^(2 −*s*)^5^ for 0.5 ≤ *s* ≤ 2. The mean of the next generation’s initial size given the current generation’s is ⟨*a*_1_|*a*_0_⟩ = *µ*(*a*_0_) = *αa*_0_ + (1− *α*), ensuring that the homeostatic mean setpoint is 1 and allowing easy comparison between different *α*’s. **(a**,**b)** *α* = 0.5. *α* satisfies the necessary and sufficient condition for homeostasis (*α* ≤1*/s*_*max*_), thus all moments converge to their homeostatic limits. (b) Inset in (B) is a zoomed plot to show rapid convergence. **(c**,**d)** *α* = 0.972. The general condition for homeostasis is not satisfied and higher moments do not converge. However, since 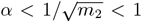, the second moment and mean converge, showing partial attainment of homeostasis in this case. **(e**,**f)** *α* = 0.973. The value of *α* is greater than 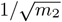, thus variance does not converge. However, since *α <* 1, the mean still converges. **(g**,**h)** *α* = 1. Even the condition for *a*_0_-independent convergence of mean is no longer satisfied, resulting in complete breakdown of homeostasis. In summary, these plots demonstrate that as *α* increases, the rates of convergence of moments slow down, and as it crosses 1*/s*_max_ followed by the limits (*m*_*k*_)^*−*1*/k*^, determined by the moments *m*_*k*_ of Π as derived in the main text, the corresponding moments no longer converge and homeostasis is broken.

### E Restrictions on the mean rescaled distribution Π(*s*) and the slope of the calibration curve *α*

We recapitulate the general necessary and sufficient homeostasis condition previously derived:

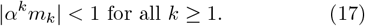

(Recall that for *α <* 0 we also require 0 *< a*_0_ *< β/*|*α*|.) This relation places strong constraints on the meanrescaled distribution Π(*s*) and its relationship with *α*. We first show that Π(*s*) must have finite support. In other words, the mean-rescaled intial size of the current generation given the initial size of the previous generation must have a maximum allowed value. To demonstrate this, we proceed as follows. Consider the case where Π(*s*) has infinite support. For *any* nonzero value *α* = *α*_0_,

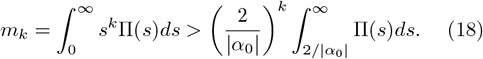

Denoting the integral on the right hand side of the inequality sign by ℐ,

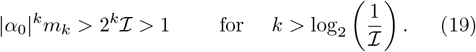

Since the preceding condition on *k* can always be satisfied because the infinite support for Π implies ℐ*>* 0, we conclude that all homeostasis conditions, Eq. (17), cannot be satisfied for any arbitrary non-zero value of *α*, however small. *Hence we conclude that* Π(*s*) *needs to have finite support if cell size homeostasis is to be achieved and sustained*. In turn, the implication is that the support of Π(*s*) extends to a maximum value, *s*_max_.

We now proceed to determine the allowed values of *α* that satisfy the homeostasis conditions Eq. (17), when Π(*s*) has finite support. Since

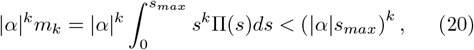

the homeostatis conditions |*α*|^*k*^*m*_*k*_ *<* 1 will always be satisfied for |*α*| ≤ 1*/s*_max_ (provided Π is not a Dirac delta function, in which case the necessary and sufficient condition for homeostasis simply reduces to *α <* 1). In the complementary scenario when |*α*| *>* 1*/s*_max_, consider any value *b* satisfying 1 *< b <* |*α*|*s*_max_. Then,

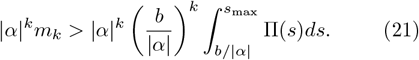

Denoting by ℐ the integral on the right hand side of the inequality,

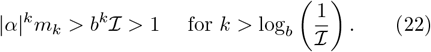

Since the condition on *k* can always be satisfied by large enough values of *k*, all homeostasis conditions cannot hold for |*α*| *>* 1*/s*_max_. Combining the above proofs that the homeostasis conditions Eq. (17) can always be satisfied when |*α*| ≤ 1*/s*_max_, and a subset of those conditions will always be violated when |*α*| *>* 1*/s*_max_, we have derived a remarkable condition:

Cell size homeostasis is obtained for all *α* satisfying |*α*| ≤ 1*/s*_max_, where *s*_max_ is the upper limit of the support of the universal meanrescaled distribution, Π(*s*), independent of other details of the shape of Π(*s*). When *α <* 0 we also require 0 *< a*_0_ *< β/*|*α*|.

To gain physical insight into this simple limit on *α*, we note that it can be rationalized by requiring that the coefficient of *a*_0_ in the expression for *a*_*n*_, Eq. (7), decrease as *n* → ∞. Since increasing *n* by one introduces one extra factor of the form *αs*, which can have a maximum value of *αs*_max_, the coefficient would be non-increasing as *n* increases, as long as *αs*_max_ ≤1. But provided the distribution of *s* is not a Dirac delta, as *n* grows large, the probability of choosing the maximum value of *s* each draw decreases to zero, ensuring the coefficient of *a*_0_ decreases. The preceding heuristic argument is also rigorously borne out by our exact probabilistic analysis. If Π is a Dirac delta function, the only possible value of *s* is 1, and the condition for homeostasis is *α <* 1, which again ensures that the coefficient of *a*_0_ decreases over successive generations.

Fig. 4 A shows a comparison between the experimentally measured values of ≤1*/α* and *s*_max_ in *E. coli* and *B. subtilis*. From this comparison it is clear that our predicted general condition for homeostasis, *s*_max_ ≤ 1*/α*, is satisfied by experimentally observed *E. coli* and *B. subtilis* growth and division dynamics, under the growth conditions shown.

To explicate and visualize the breakdown of homeostasis as the theoretical condition |*α*| ≤ 1*/s*_max_ is violated, we take advantage of numerical simulations based on the theoretical framework proposed here. In Fig. 5 we show the simulated distributions of initial sizes over successive generations for a range of *α* values for hypothetical cells. These include cases for which the condition for homeostasis is not satisfied. Fig. 6 shows the generational evolution of the first two moments of cell size, as *α* passes through different homeostasis thresholds. When |*α*|*<* 1*/s*_max_, the cell size distribution converges to a well-defined distribution with a short tail. When 1 *>* |*α*| *>* 1*/s*_max_, the mean and some lower moments converge to values independent of the starting generation size *a*_0_, but higher moments do not, serving as indicators of the expected breakdown of homeostasis. This breakdown is reflected in the distributions developing long tails after sufficiently many generations have elapsed. Finally, when *α >* 1, we predict that even the mean cannot attain an *a*_0_-independent value (Fig. 6 G); in confirmation, the distributions become qualitatively different for distinct *a*_0_ values, showing a spectacular breakdown of homeostasis (see Fig. 5).

## IV. CONCLUDING REMARKS

In summary, we have developed a principled data-informed stochastic theoretical framework to characterize cell size homeostasis of bacterial cells in balanced growth conditions. Prior works typically describe homeostasis building on a quasi-deterministic scheme, with noise “added on top” motivated by analytical or computational tractability [22, 26–29]. The condition for homeostasis obtained from quasi-deterministic heuristics, *α <* 1, needs to be supplemented when the fully stochastic picture is considered. In this work we have shown that the necessary and sufficient condition for homeostasis, when considering the data-informed [1] inherently stochastic formulation of the problem, is |*α*| ≤ 1*/s*_max_, where *s*_max_ *>* 1 is the upper limit of the support of the mean-rescaled distribution Π, introduced in Eq. (3). By reanalyzing published experimental data in *E. coli* and *B. subtilis* (Figs. 2 and 3 respectively), we have not only shown compelling data-theory matches, but also that this new homeostatis condition, |*α*| ≤ 1*/s*_max_, is indeed satisfied by these data (see downward pointing arrows in Fig. 4).

In addition to this condition requiring all moments of the steady state initial size distribution to be finite, we have also derived the conditions for only specific moments to be finite. An alternate condition for all moments to be finite was previously derived for a linear model with additive noise in [20]. That said, this work also obtained a constraint on *α* identical to the deterministic models (*α <* 1), while experimental evidence shows *α* indeed lies very close to (and below) our derived limit of 1*/s*_max_ (Fig. 4).

For the growth conditions reported here and in [1], the necessary and sufficient condition for cell size homeostasis appears to be satisfied. But under starvation or other stressful growth conditions, the constraints on higher moments may not be satisfied, because in practice, occurrence of rare instances of instability, despite reducing the overall population fitness, may be acceptable from the standpoint of overall population survival. This can happen when only the higher moments diverge, but not lower moments such as the mean and variance, leading to a fraction of the population having runaway sizes (perhaps through the phenomenon of filamentation, observed primarily in stressful growth conditions). We have further analyzed the consequences of these constraints not being satisfied in [30].

While the theoretical framework of intergenerational size dynamics presented here is data-informed and accurately describes the kinematics of cell size homeostasis, it is mechanistically agnostic. In complementary work [31] we have addressed possible architectural underpinnings of the observed intergenerational scaling law leading to these precision kinematics.

Our theoretical framework is built on two experimentally-observed emergent simplicities [1]: first, the inter-generational initial size dynamics are Markovian, and second, the conditional distribution of the next generation’s initial size, conditioned on the current generation’s initial size, when rescaled by its mean value, results in a distribution (Π) that is invariant of this generation’s initial size. Since the mean-rescaled distribution changes from growth condition to growth condition, even for the same organism (as shown in [1]), how these results generalize to time-varying growth conditions remains to be seen.

## AUTHOR CONTRIBUTIONS

K.J., R.R.B., and S.I.-B. conceived of and designed the research; K.J. developed the theoretical framework under the guidance of S.I.-B.; K.J., R.R.B. and S.I.B. performed analytic calculations; C.S.W. curated the datasets; K.J. and C.S.W. performed data analyses; K.J. performed simulations under the guidance of R.R.B. and S.I.-B.; K.J., C.S. W., R.R.B., and S.I.-B. wrote and revised the paper; S.I.-B. supervised the research.

## ACKNOWLEDGEMENTS

R.R.B and S.I.-B gratefully acknowledge Purdue University Startup funds and the Purdue Research Foundation for supporting the research and the collaboration. S.I.-B. thanks the Purdue College of Science Dean’s Special Fund, and the Showalter Trust for financial support. K.J. and S.I.-B. acknowledge support from the Ross-Lynn Fellowship award. We are grateful to Sattar Taheri Araghi, Suckjoon Jun for generously sharing the previously published single-cell datasets that are utilized in this study, and to Sattar Taheri Araghi, Suckjoon Jun, and Fangwei Si for kindly explaining the details of their methodology and analysis.

### APPENDIX: ANALYTICAL METHODS

#### A. Homeostasis condition for general raw moment of *a*_*n*_

Before calculating the general raw moments of *a*_*n*_, we convert to rescaled variables 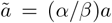 and 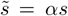. Thus, 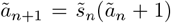. The random variable 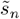 has moments 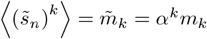. Note the special cases: 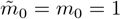 and 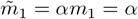.

Using these rescaled variables, the appropriate stochastic relation Eq. (7) from the main text becomes:

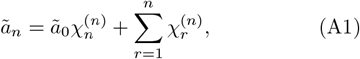

where,

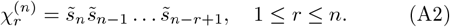

Correlations of the newly-defined random variables 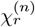 have the following values in terms of the moments of 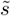:

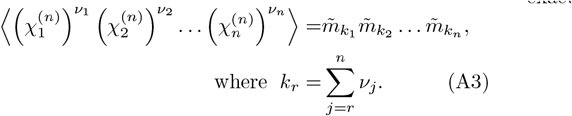

In the above expression, all *ν*_*j*_ 0 and 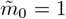. Raising Eq. (A1) to the *k*^th^ power and taking average, we find an exact expression for the *k*^th^ moment of ã_*n*_:

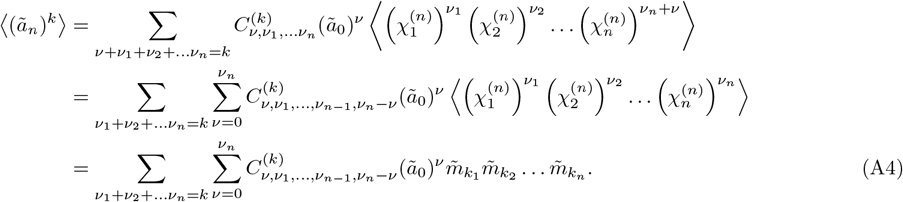

(Using ⟨ (*a*_*n*_)^*k*^⟩ = (*β/α*)^*k*^ ⟨(ã_*n*_)^*k*^*⟩* with the above equation yields an exact expression for the *k*^th^ moment of *a*_*n*_.) Here, all sums are over non-negative values of (*ν, ν*_1_, *…, ν*_*n*−1_, *ν*_*n*_),

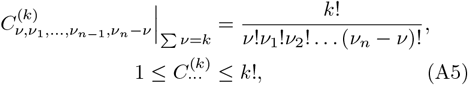

and we have used Eq. (A3) in the last step. Noting that *k*_*n*_ = *ν*_*n*_, can further simplify Eq. (A4):

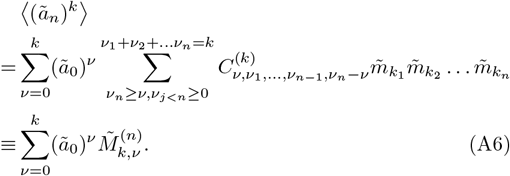

To aid in the upcoming analyses, we define a new quantity 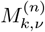:

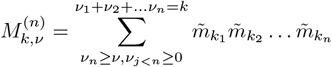

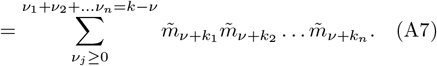

Before finding the asymptotic limits of 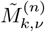 as *n* → ∞, we first consider the asymptotic limits of 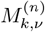.

Considering the last expression in Eq. (A7) and the definitions of *k*_*r*_’s given in (A3), we note that the *k*_*r*_’s form a non-increasing series in *r* bounded by *k* −*ν* above and 0 below: *k*_1_(= *k* −*ν*) ≥ *k*_2_ ≥*…*≥ *k*_*n*_(= *ν*_*n*_) ≥ 0. Also, every set of {*k*_*r*_} satisfying these in/equalities are in one-to-one correspondence with an allowed set *ν*_*j*_ in the last step in Eq. (A7). Furthermore, such a set of *k*_*r*_’s (and thus *ν*_*j*_’s) are in one-to-one correspondence with the set 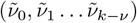, where 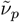 is the number of *k*_*r*_’s which are equal to *p*, 0 ≤ *p* ≤ *k* −*ν*, and 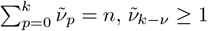 and 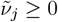 for 0 ≤ *j < k* − *ν*. Thus, the sum over *{ν*_*j*_*}* in Eq. (A7) can be replaced by a sum over 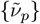, with the substitution 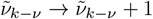:

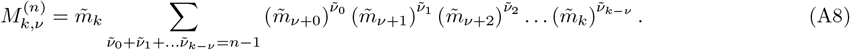

First, consider the case *ν* = 0, which controls the value of the ã_0_-independent term in Eq. (A6),

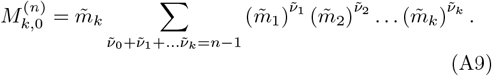

Noting that 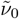 does not appear in the sum at all, the sum over 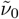 can be performed immediately, changing the sum over the remaining 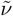’s as follows:

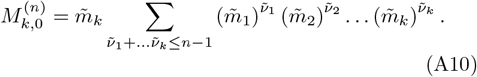

As *n* → ∞,

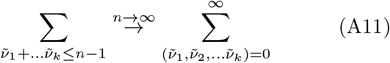

and so

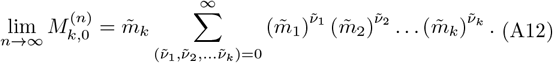

The sums converge only when 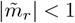 for *r* = 1, 2, *…, k*, when

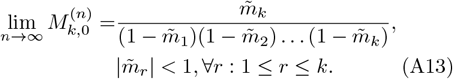

Meanwhile, for 0 *< ν* ≤ *k*, Eq. (A8) becomes

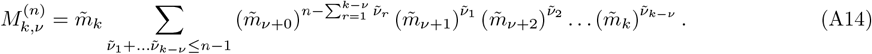

The conditions for convergence of *M*_*k*,0_ also ensure the convergence of the sum above, since

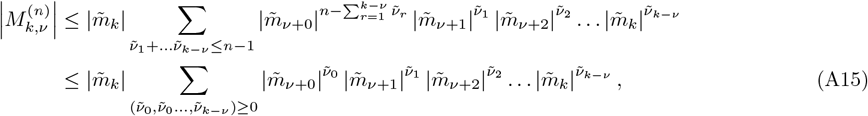

and the last expression converges when the conditions in Eq. (A13) are satisfied. Moreover, setting 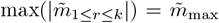 with 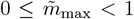 (condition of convergence of *M*_*k*,0_), from Eq. (A14),

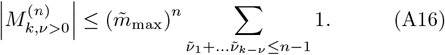

As *n* → ∞, the sum grows as a power law of *n* with a fixed exponent 𝒪 (*k*), while the outer factor vanishes exponentially with *n*. Thus, as *n* → ∞, the expression on the right vanishes and so:

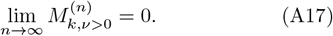

*1. Homeostasis condition for non-negative α values*

For non-negative *α* values, using Eq.s (A5), (A6), and (A7),

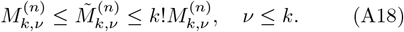

Thus, as *n* → ∞, the convergence and possible range of asymptotic values of each coefficient 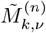 is ensured by the convergence and asymptotic values for the corresponding 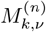.

Consequently, the conditions that guarantee the convergence of the ã_0_-independent part of (ã_*n*_)^*k*^ also ensure that all initial generation size dependence vanishes as *n* → ∞. To summarize, using Eqs. (A13), (A17) and (A18) in Eq. (A14), we find:

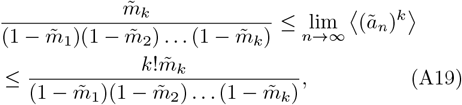

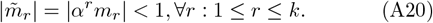

Rewriting Eq. (A19) in terms of the original size variables (using the relation *m*_1_ = 1), we summarize the final result of this analysis for *α* ≥ 0:

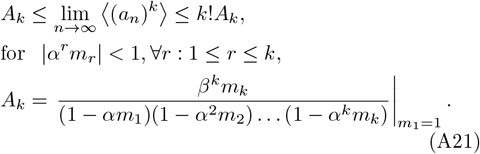

##### 2. Homeostasis condition for negative α values

We now find the homeostasis condition for negative values of *α*, when we need to consider an additional complexity: the size cannot become negative. In the main text we have showed that the conditions for homeostasis when *α* ≥ 0: |*α*^*k*^*m*_*k*_| *<* 1 for all *k* = 1, 2, …, are equivalent to the single condition |*α*| *<* 1*/s*_max_, where *s*_max_ is the upper limit of the range of Π(*s*). We will now show that in addition to this condition, the requirement 0 *< a*_0_ *< β/* |*α*| needs to be met for homeostasis to occur when *α <* 0.

We take a slightly different approach from that in the previous section, and use the stochastic map *a*_*n*+1_ = *s*_*n*_(*αa*_*n*_ + *β*) corresponding to Eq. (5). We consider *s* to be restricted between *s*_*min*_ and *s*_*max*_, the two limits of the support for Π(*s*).

First, consider the special case *s*_*min*_ = *s*_*max*_(= 1). This makes Π a Dirac delta function, and *a*_*n*+1_ = *αa*_*n*_ + *β*. The solution for *a*_*n*_ is given by Eq. (8), where ⟨*a*_*n*_⟩ = *a*_*n*_ for this case. Thus, |*α*| *<* 1 is necessary and sufficient to ensure convergence as *n* → ∞, corresponding to the homeostasis requirement for the *α >* 0 case. Additionally however, we also require all *a*_*n*_ *>* 0. Considering Eq. (8) again, since the magnitude of the *α*^*n*^-containing second term decreases as *n* increases, and the first term is positive, it is enough to find the condition for which *a*_*n*_ *>* 0 for the lowest even and odd *n*. For *n* = 0, this requirement is trivially true, since *a*_0_ *>* 0. For *n* = 1, we find the condition *a*_0_ *< β/* |*α*|. Thus, the necessary and sufficient condition for homeostasis and guaranteed positive initial size is |*α*| *<* 1 and *a*_0_ *< β/*|*α*|.

Next, consider the remaining general case, *s*_*min*_ *< s*_*max*_. Here,

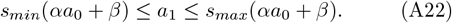

For the next generation,

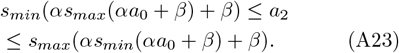

Continuing further,

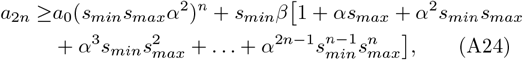

etc. Simplifying, we find,

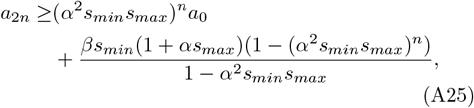

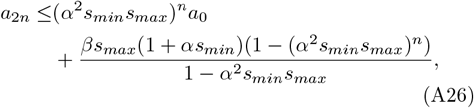

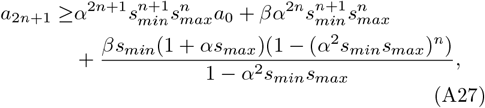

and finally,

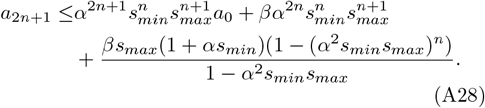

Simplifying further,

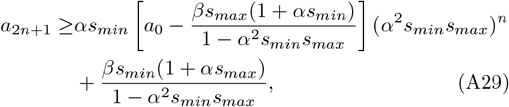

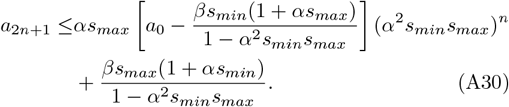

Since *α* is negative, but *a*_1_ and *a*_2_ are positive,

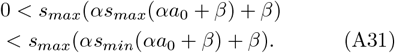

Continuing like this, for every non-negative *n*, we need

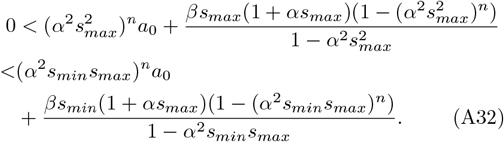

If |*αs*_*max*_| *>* 1, 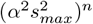 diverges faster than (*α*^2^*s*_*min*_*s*_*max*_)^*n*^ for large *n*, which violates the above inequality. Thus, we arrive at a necessary condition for homeostasis, which is the same as that for the *α* ≥ 0 case:

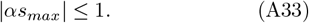

If this holds, then it is clear that both limits in Eq.s A25 and A26 are positive and the lower limit is less than or equal to the upper limit. Now we just need to check Eq.s A29 and A30. In these equations, since (*α*^2^*s*_*min*_*s*_*max*_)^*n*^ decreases with *n*, and the remaining term is positive, as long as the limits are positive for the lowest *n*, they will be positive for all *n*. This is true for *n* = 0 if, and only if, *a*_0_ *< β/* |*α*|. Thus, the limits are positive under this condition. It now remains to check that the lower limit is less than or equal to the upper limit. Subtracting the lower limit from upper limit,

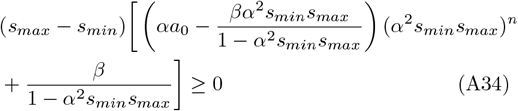

Again, since (*α*^2^*s*_*min*_*s*_*max*_)^*n*^ decreases with *n*, and the remaining term is positive, as long as the above expression is non-negative for the lowest *n*, it will be non-negative for all *n*. Checking for *n* = 0, we again obtain the condition: *a*_0_ ≤ *β/*|*α*|.

Thus, when |*α*| ≤ 1*/s*_*max*_, *a*_0_ *< β/* |*α*| is the necessary and sufficient condition for initial size to be restricted to positive values when *α* is negative.

We now proceed to determine the sufficiency of the above conditions for convergence of the moments. From Eqs (A5), (A6), and (A7), we have,

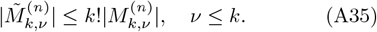

Now, as we have derived in the main text, |*α*| ≤ 1*/s*_*max*_ ensures that |*α*^*k*^*m*_*k*_| *<* 1 for all *k*. This in turn ensures Eq.s (A13) and (A17) hold. Thus, from Eq.s (A6), (A35), (A13), and (A17),

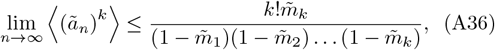

or, in terms of the original variables,

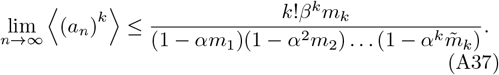

Thus, the moments converge, showing that |*α*| ≤ 1*/s*_*max*_ is sufficient. Thus, for negative *α* values, we have proved the necessary and sufficient conditions for homeostasis: |*α*| ≤ 1*/s*_*max*_ and *a*_0_ *< β/*|*α*|.

#### B. Exact solution for the asymptotic limits of the moments of *a*_*n*_

To find the exact solution for the asymptotic limits of the moments, we start by noting that as *n* → ∞, only the *ν* = 0 term survives in Eq. (A4) as long as the condition for homeostasis is satisfied (because the result becomes independent of *ã*_0_, as derived in previous sections). Thus, replacing the value of *C* from Eq. (A5),

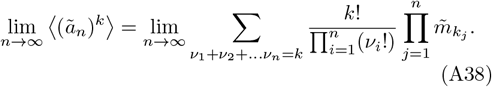

Now, as *n* → ∞, there will be infinitely many *v*_*i*_’s, and all of them are restricted to non-negative integer values. Since their sum is finite, only a finite number of them can take non-zero values. From the definition of *k*_*r*_ in Eq. (A3), we have *k*_1_ = *k* and *k*_*r*_ ≤ *k*_*r*+1_ ≤ 0, and there are infinitely many *k*_*r*_’s, and *v*_*r*_ = *k*_*r*_ − *k*_*r*+1_. This set of *k*_*r*_’s (and thus *ν*_*j*_’s) are in one-to-one correspondence with the set (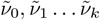), where 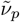 is the number of *k*_*r*_’s which are equal to *p*, 0 ≤ *p* ≤ *k*, and 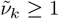, and 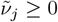 for 0 ≤ *j < k*.

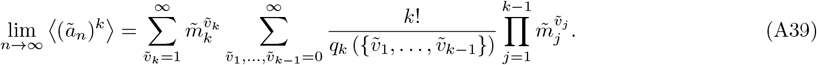

Here, the function *q* is a replacement for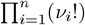. Multiplying by *v*_*i*_! will not change the value of *q* as long as *v*_*i*_ is either 0 or 1. To contribute to *q, v*_*i*_ must be greater than 1. But, *v*_*r*_ = *k*_*r*_ − *k*_*r*+1_, with *k*_*r*_ ≤ *k*_*r*+1_. Thus, if *v*_*i*_ *>* 1 and *k*_*i*_ = *p*, then, *k*_*r*+1_ *> p* + 1, i.e., 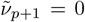. Thus, each value of *v*_*i*_ that contributes to *q* (i.e., is greater than 1), corresponds to a zero in the sequence of 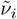’s. Also, if there is a sequence of consecutive zeros of length

*l* in the sequence of 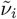, then there would be a corresponding *v*_*j*_ with value equal to *l* + 1, which contributes (*l* + 1)! to *q*. Thus, the value of *q*_*k*_ (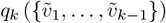) is determined by setting the value to 1, then multiplying it by (*l* + 1)! for each set of consecutive zeros of length *l* in the sequence of 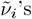 for all *l*.

Now, converting the result to the original variables of *a* and *m*,

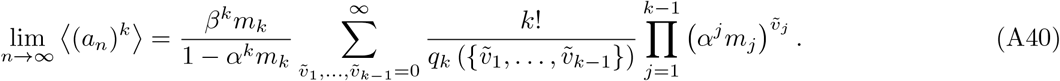

### APPENDIX: ADDITIONAL FIGURES

**Appendix Figure A1.**
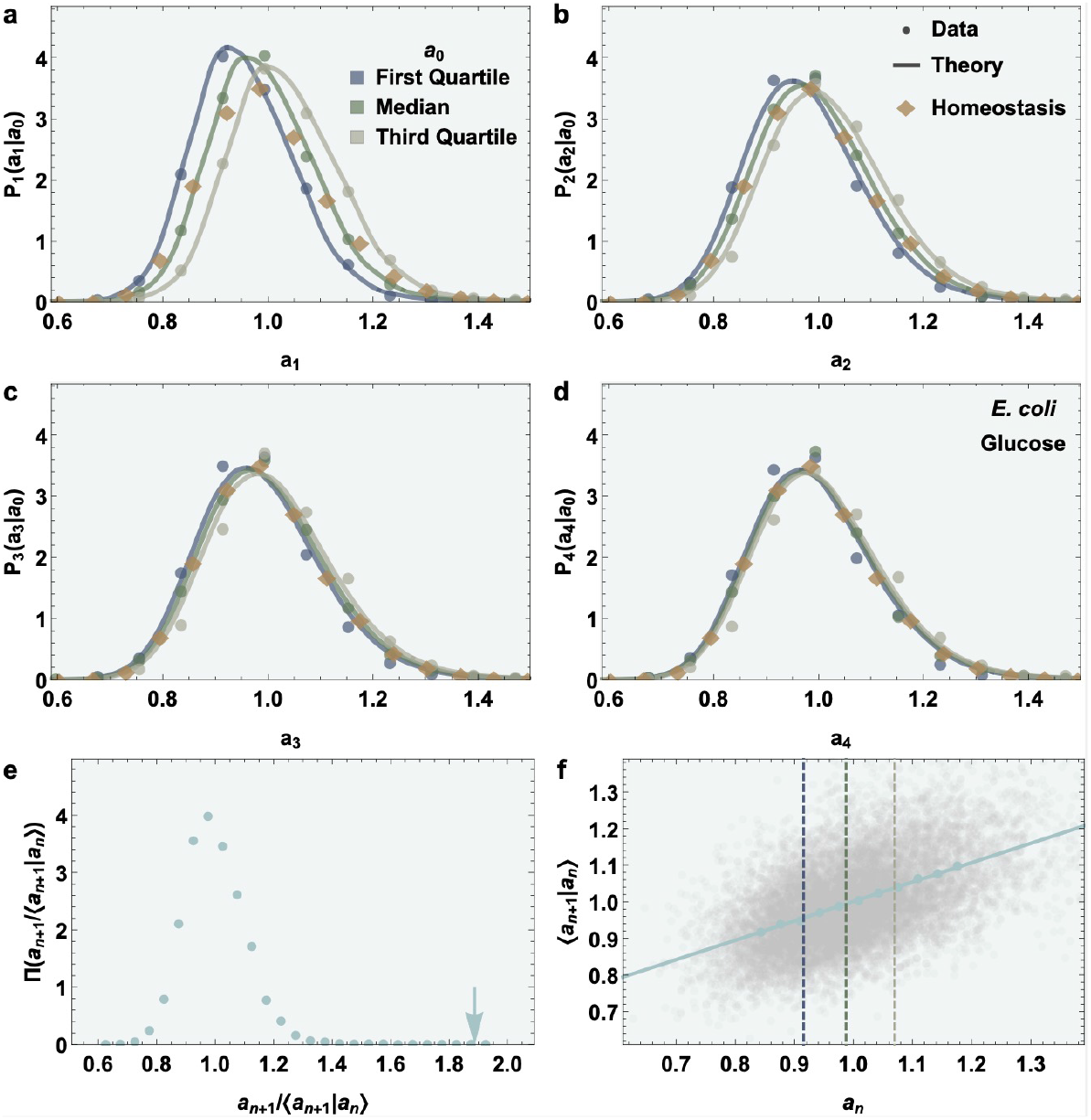
Our theoretical framework accurately predicts, with no fitting parameters, the stochastic intergenerational cell size dynamics leading to homeostasis, for *E. coli* cells. See Fig. 2 caption in the main text for further details. These plots correspond to *E. coli* cells grown in Glucose media. Cell length was used as a measure of cell sizein these datasets. See Fig. 3 in the main text for similar application to *B. subtilis* cells, and the Appendix for application to other growth conditions for both species.

**Appendix Figure A2.**
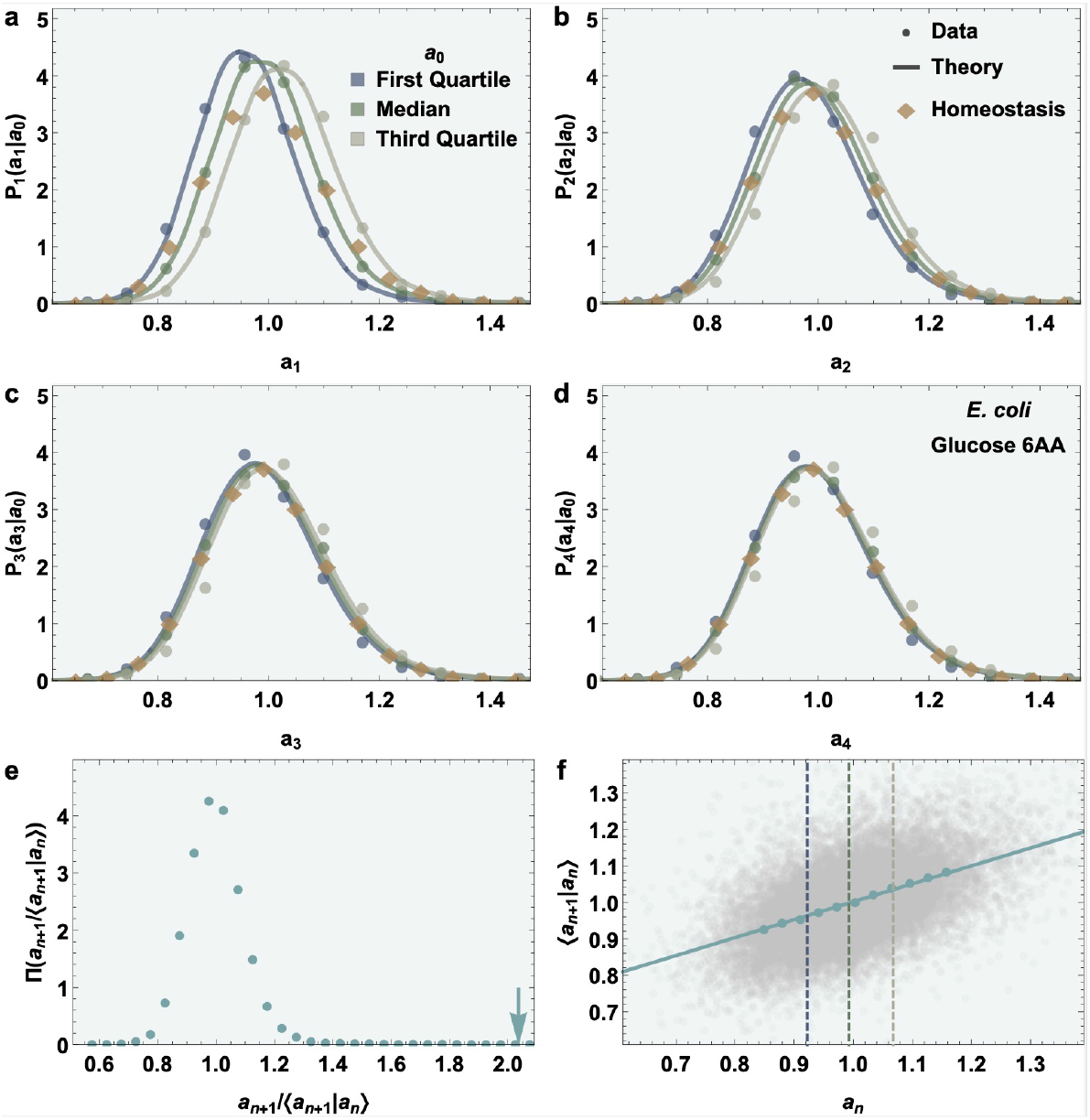
Our theoretical framework accurately predicts, with no fitting parameters, the stochastic intergenerational cell size dynamics leading to homeostasis, for *E. coli* cells. See Fig. 2 caption in the main text for further details. These plots correspond to *E. coli* cells grown in Glucose 6AA media. Cell length was used as a measure of cell size in these datasets. See Fig. 3 in the main text for similar application to *B. subtilis* cells, and the Appendix for application to other growth conditions for both species.

**Appendix Figure A3.**
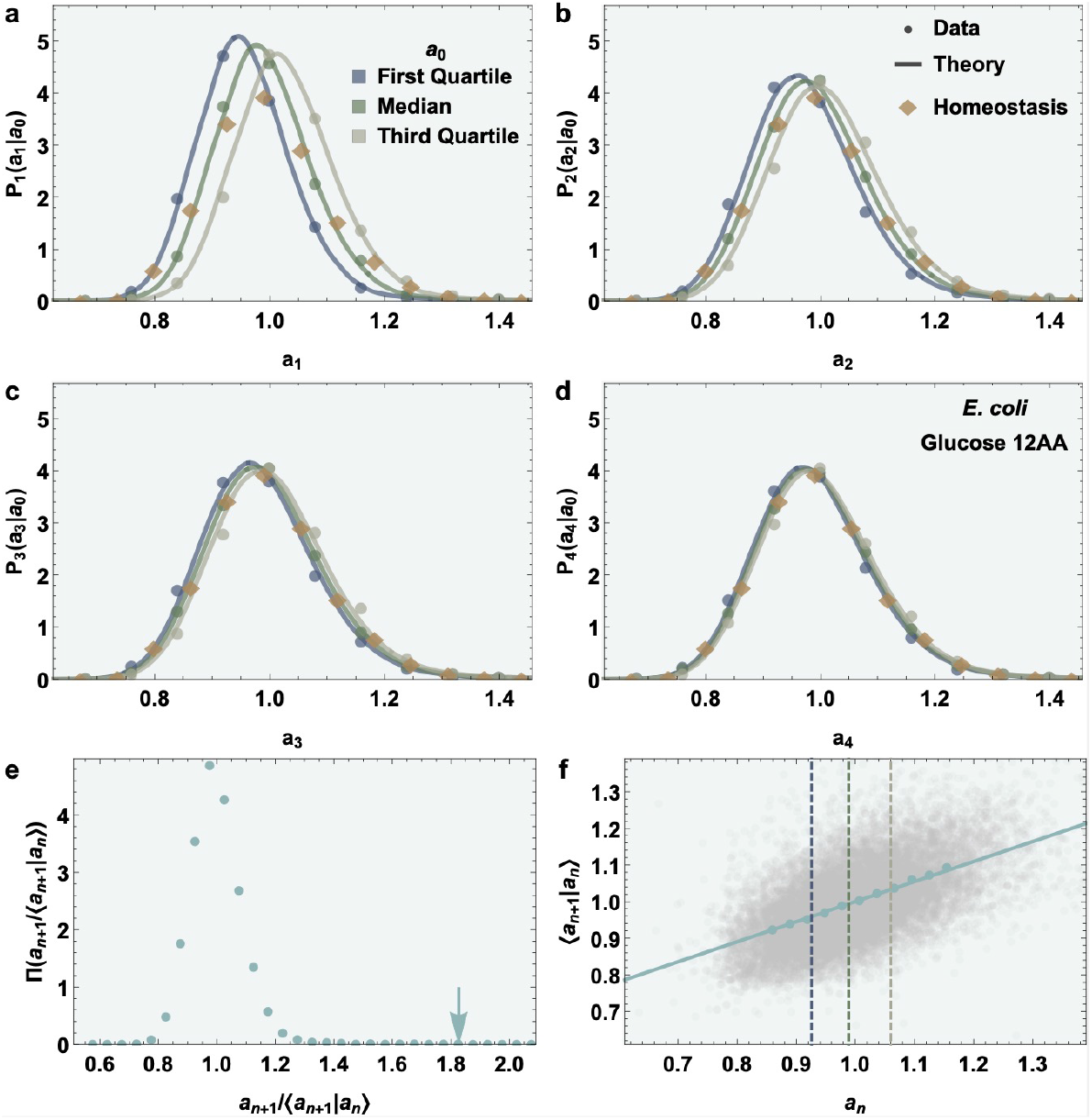
Our theoretical framework accurately predicts, with no fitting parameters, the stochastic intergenerational cell size dynamics leading to homeostasis, for *E. coli* cells. See Fig. 2 caption in the main text for further details. These plots correspond to *E. coli* cells grown in Glucose 12AA media. Cell length was used as a measure of cell size in these datasets. See Fig. 3 in the main text for similar application to *B. subtilis* cells, and the Appendix for application to other growth conditions for both species.

**Appendix Figure A4.**
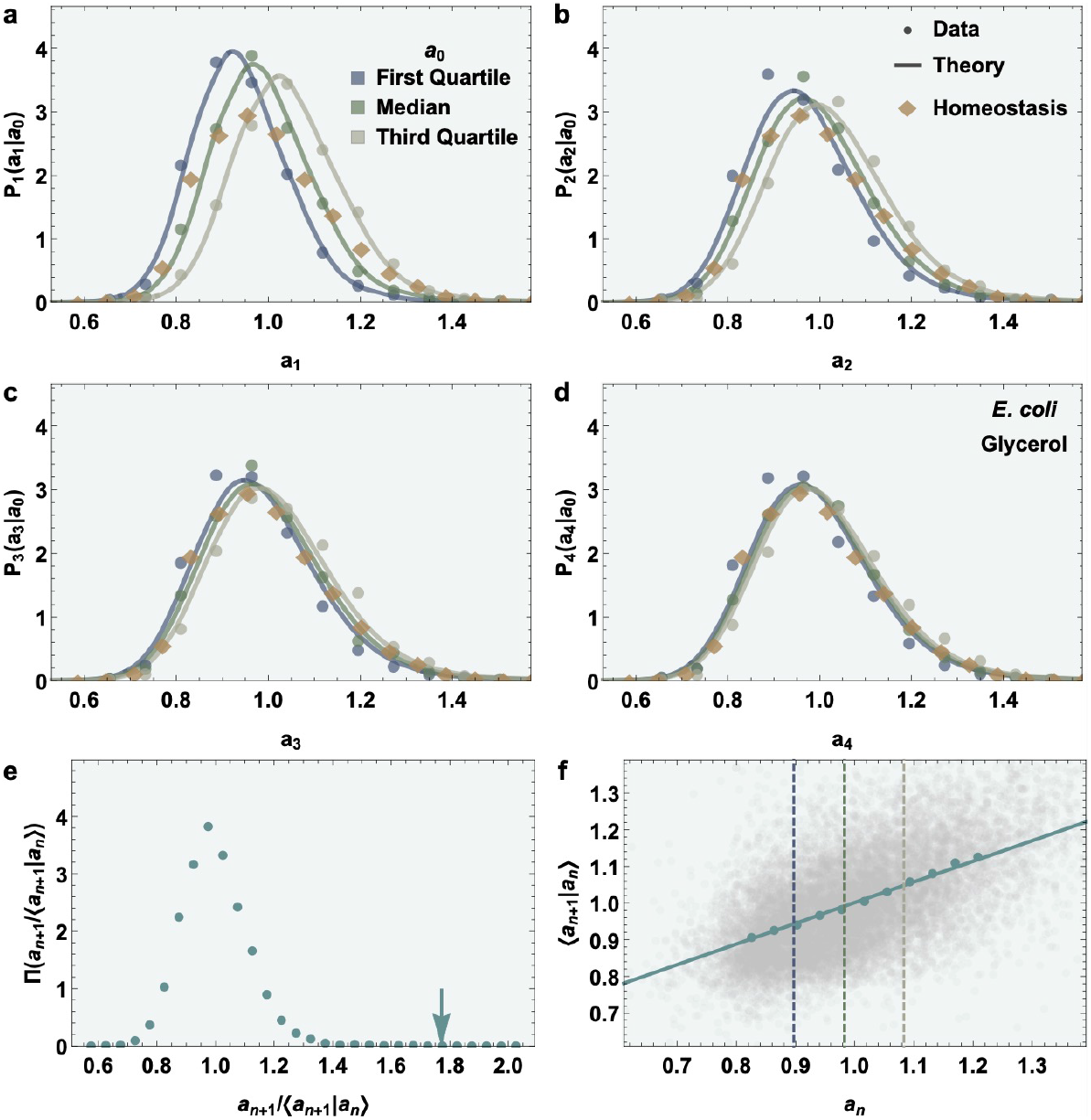
Our theoretical framework accurately predicts, with no fitting parameters, the stochastic intergenerational cell size dynamics leading to homeostasis, for *E. coli* cells. See Fig. 2 caption in the main text for further details. These plots correspond to *E. coli* cells grown in Glycerol media. Cell length was used as a measure of cell size in these datasets. See Fig. 3 in the main text for similar application to *B. subtilis* cells, and the Appendix for application to other growth conditions for both species.

**Appendix Figure A5.**
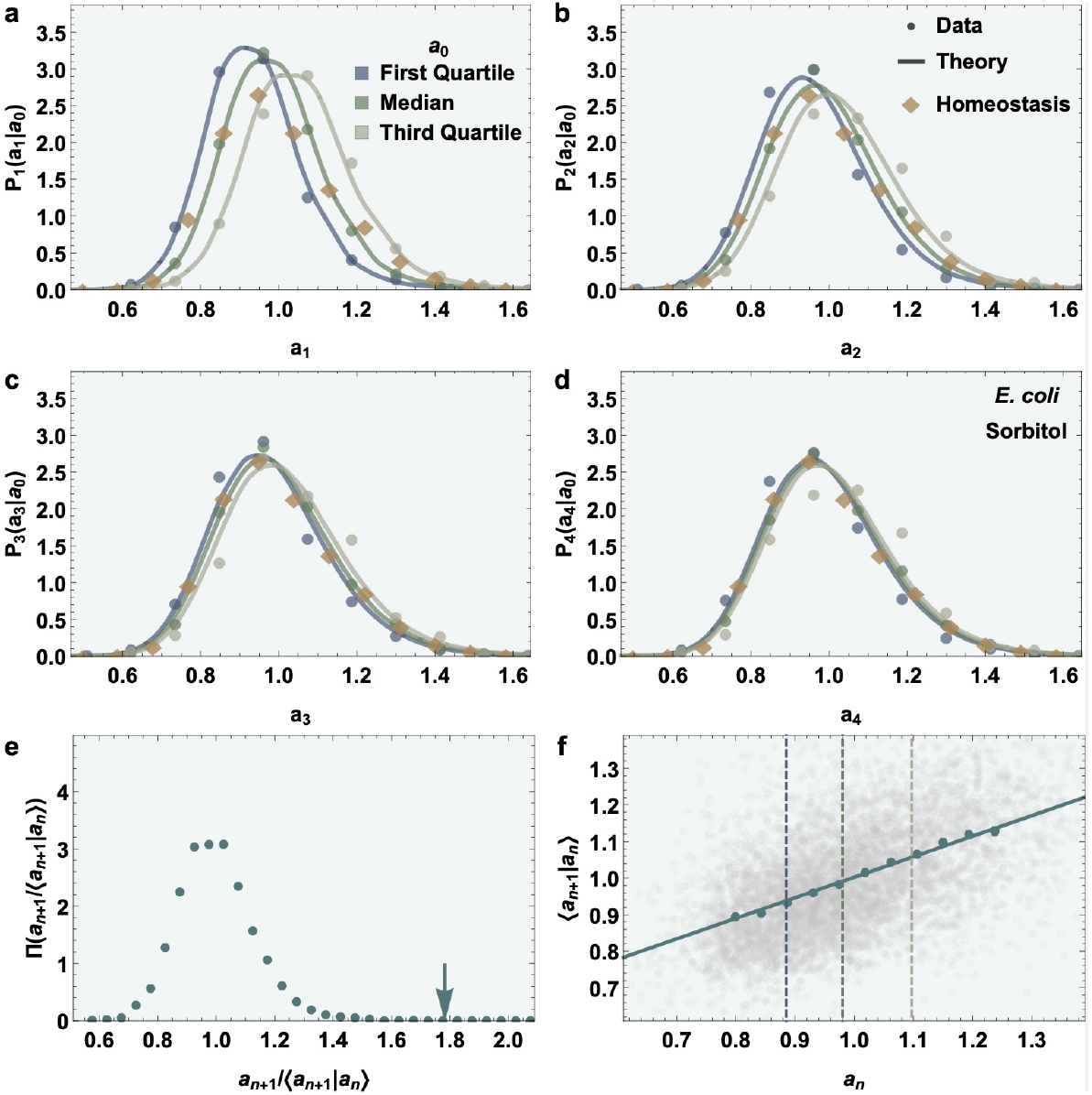
Our theoretical framework accurately predicts, with no fitting parameters, the stochastic intergenerational cell size dynamics leading to homeostasis, for *E. coli* cells. See Fig. 2 caption in the main text for further details. These plots correspond to *E. coli* cells grown in Sorbitol media. Cell length was used as a measure of cell size in these datasets. See Fig. 3 in the main text for similar application to *B. subtilis* cells, and the Appendix for application to other growth conditions for both species.

**Appendix Figure A6.**
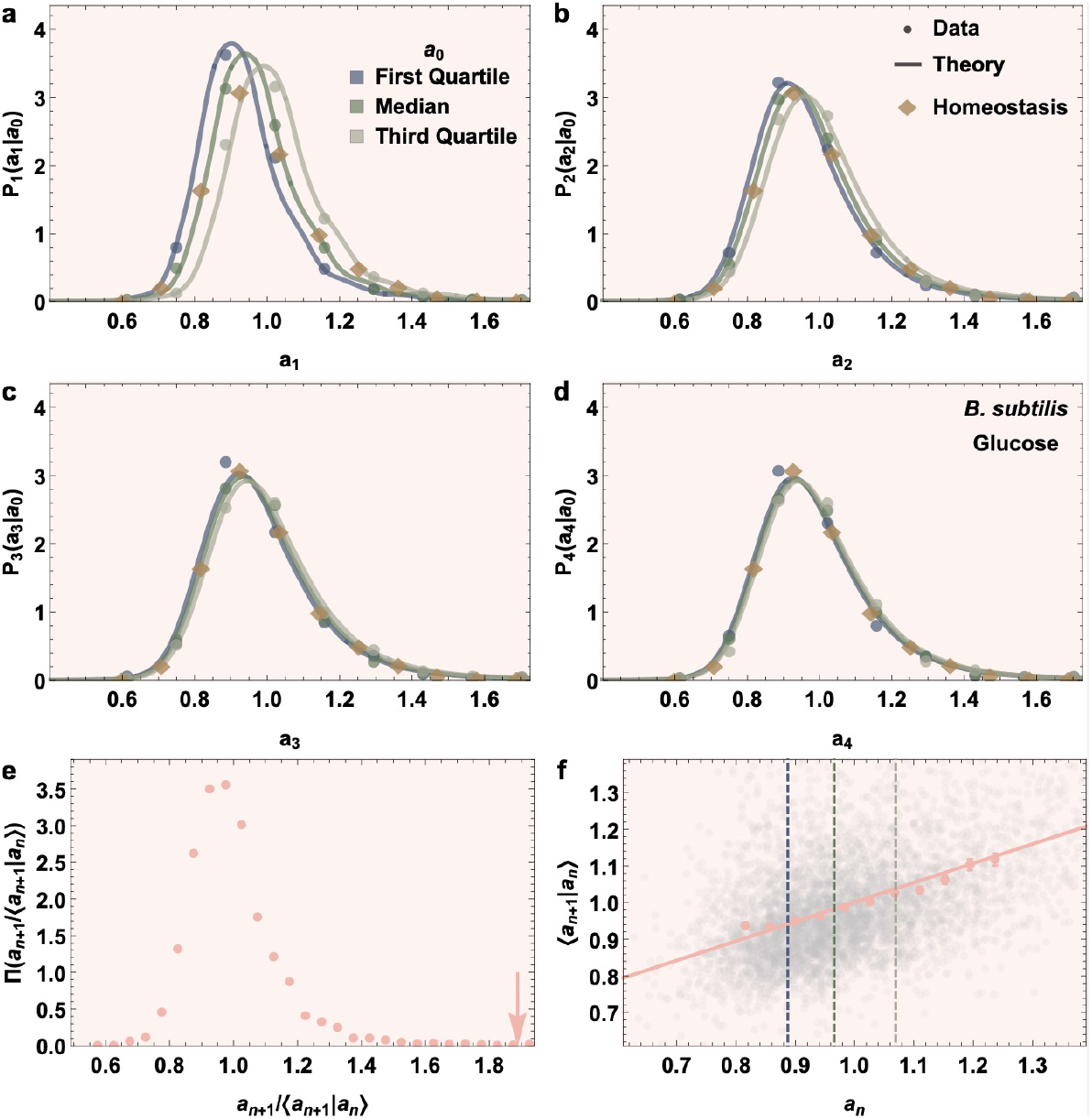
Our theoretical framework accurately predicts, with no fitting parameters, the stochastic intergenerational cell size dynamics leading to homeostasis, for *B. subtilis* cells. See Fig. 3 caption in the main text for further details. These plots correspond to *B. subtilis* cells grown in Glucose media. Cell length was used as a measure of cell size in these datasets. See Fig. 2 in the main text for similar application to *E. coli* cells, and the Appendix for application to other growth conditions for both species.

**Appendix Figure A7.**
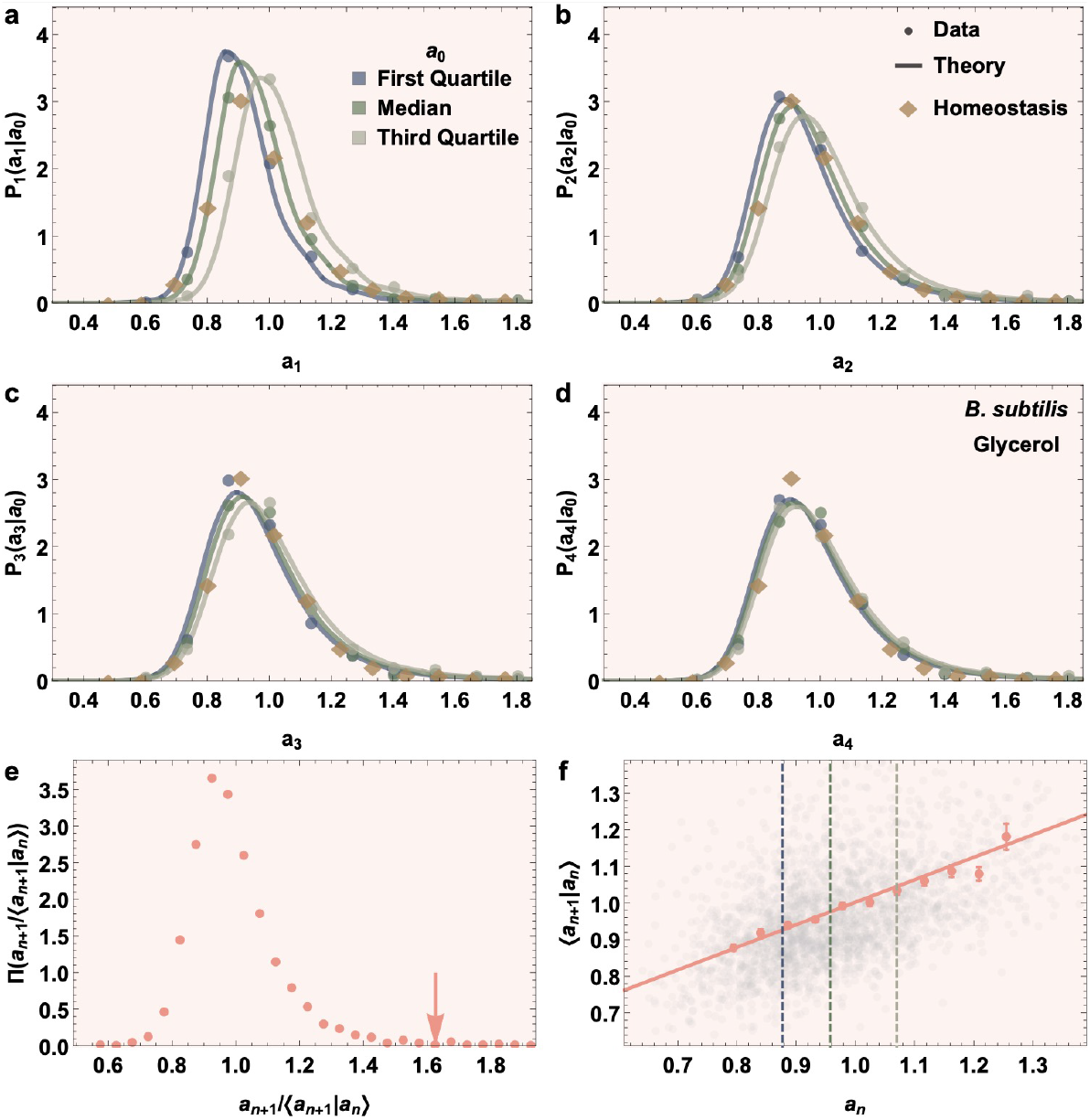
Our theoretical framework accurately predicts, with no fitting parameters, the stochastic intergenerational cell size dynamics leading to homeostasis, for *B. subtilis* cells. See Fig. 3 caption in the main text for further details. These plots correspond to *B. subtilis* cells grown in Glycerol media. Cell length was used as a measure of cell size in these datasets. See Fig. 2 in the main text for similar application to *E. coli* cells, and the Appendix for application to other growth conditions for both species.

**Appendix Figure A8.**
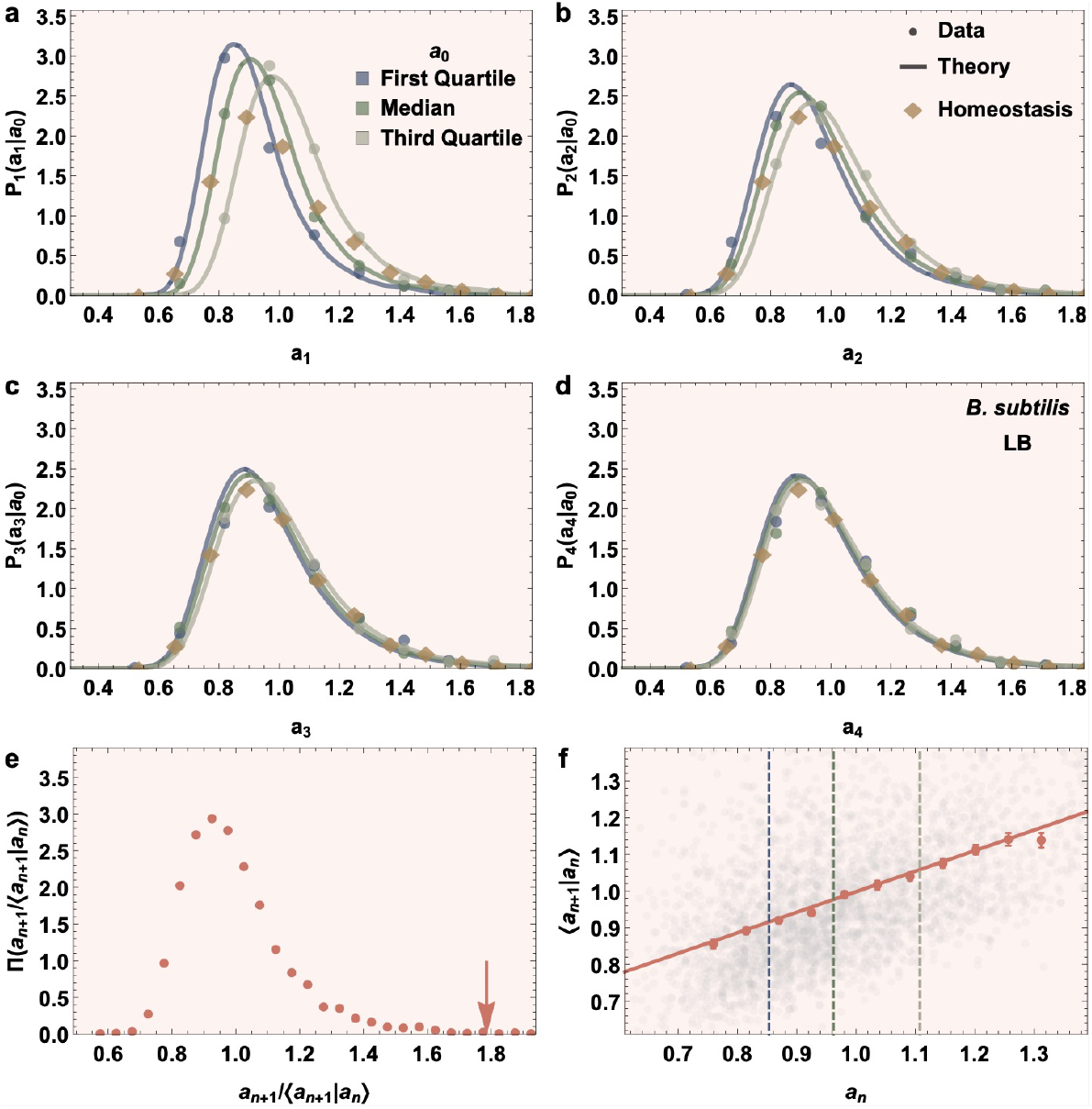
Our theoretical framework accurately predicts, with no fitting parameters, the stochastic intergenerational cell size dynamics leading to homeostasis, for *B. subtilis* cells. See Fig. 3 caption in the main text for further details. These plots correspond to *B. subtilis* cells grown in LB media. Cell length was used as a measure of cell size in these datasets. See Fig. 2 in the main text for similar application to *E. coli* cells, and the Appendix for application to other growth conditions for both species.

